# Inhomogeneous Tau polymerization, core–shell organization, and seed formation during Tau condensate aging

**DOI:** 10.64898/2026.03.18.711671

**Authors:** Maximilian Franck, Abin Biswas, Pin-Lian Jiang, Martin Fernandez-Campo, Satabdee Mohapatra, Rithika Sankar, Alvaro Dominguez-Baquero, Leandre Ravatt, Boglarka K. Nagy-Herczeg, Ozan Kantar, Janine Hochmair, Thorsten Mielke, Lisa Diez, Fan Liu, Michael Krieg, Simone Reber, Susanne Wegmann

## Abstract

Several proteins aggregating in neurodegenerative diseases spontaneously segregate into liquid condensates, which can catalyze protein aggregation. How liquid-solid transitions are catalyzed in the confined condensate volume is not clear. For the microtubule associated protein Tau, aggregating intra-neuronally in Alzheimer’s disease, we show that, during maturation, Tau condensates convert into elastic protein networks, accompanied by inhomogeneous polymerization of the condensate interior and the formation of high-density nodes and a “shell”. During condensation, Tau molecules extend, favoring intermolecular interactions and priming for progressive parallel Tau arrangement that can enable amyloid-like Tau oligomerization. In cells, aged condensates seed small Tau clusters in cytosol and at the nuclear envelope, precursors of larger aggregates. By bridging molecular to condensate level, we present mechanistic insight into how Tau condensates evolve into pathological, beta-structure containing seeds. The interior of aged Tau condensates remains accessible for smaller molecules, providing the opportunity to molecularly target Tau seed formation inside condensates.

**Teaser:** Aging Tau/RNA/PEG condensates change their viscoelasticity and protein/RNA distribution, promoting pathological Tau species.

## Introduction

The microtubule-associated protein Tau is intrinsically disordered and can take on different assembly forms, each with unique biochemical and biophysical characteristics and cellular functions. Monomers and dimers are thought to represent the soluble “native” form of Tau, whereas oligomers are assumed to be neurotoxic (*1*), template the misfolding of naïve Tau molecules (seeding), and can mediate the spread of Tau pathology between neurons in Alzheimer’s disease (AD) (*2*). Cross beta-sheet containing Tau aggregates are the stable end product of Tau aggregation and are deposited as amyloid-like, fibrillar Tau aggregates in neuronal inclusions in the brain of patients with AD and other tauopathies (*3–5*) (*6*). The core of pathological Tau fibrils is built from densely packed parallel stacks of Tau repeat domains (TauRDs), whereby the unstructured N-terminal half (∼240 aa) and the C-terminal end (∼40 aa) of Tau remain unstructured and protrude as a polymer brush-like from the fibrillar aggregate core (*5*, *7*, *8*). This arrangement of Tau is favored by interactions of two hexapeptide motifs (PHF* and PHF) in the beginning of repeats R2 and R3 in TauRD (*9*). The molecular density (mass per length) of amyloid-like aggregated Tau was calculated as 130-160 Da/nm, corresponding to ∼3 Tau molecules (46 kDa) per nm fibril length (*10*).

Like other amyloidogenic proteins, pathological, beta-sheet containing Tau assemblies – e.g., oligomers and fibrils - can template the misfolding and aggregation of naïve Tau molecules (“seeding”), which relies on the imprinting of misfolded structure onto natively unfolded proteins of the same type. This mechanism underlies not only the progressive intra- neuronal accumulation and aggregation of Tau in disease, but it also seems to facilitate the inter-neuronal spread of Tau aggregation across neural connections. We previously showed that RNA-induced Tau condensates can template Tau aggregation *in vitro* and in cells (*11*). Condensates of RNA binding proteins that aggregate in disease, i.e., hnRNPA1 and fused in sarcoma (FUS), undergo a complex aging process, during which the formation of heterogeneous domains and networks leads to a loss of molecular mobility and a mostly elastic behavior of condensates (*12*, *13*). In cells, this type of condensate aging is suspected to catalyze pathogenic protein aggregation, e.g., in neurodegenerative diseases like amyotrophic lateral sclerosis (ALS) and frontotemporal dementia (FTD). For Tau, we previously showed that *in vitro* condensate maturation can catalyze pathological Tau seed formation (*11*, *14*). However, details of the Tau condensates maturation process – harboring valuable information to interfere with this process therapeutically - remain unclear. Unraveling the mechanism behind the transition of Tau from a liquid-condensed towards seeding-competent states will further our understanding of AD and tauopathy disease origin, progression, and etiology, and to develop strategies for therapeutic interventions attacking early phases of the Tau aggregation cascade in the brain.

To understand the process leading to the formation of seeding-competent Tau during condensate maturation, we characterized the viscoelastic phase transitions and molecular rearrangements occurring in freshly formed versus “aged” Tau/RNA/PEG condensates. We find that the aging process is characterized by substantial hardening of the condensates, which can be attributed to the progressive dynamic arrest of Tau molecules. We observe network-like polymerization and the formation of densely packed domains/structures – notably, lacking beta-structure - of Tau inside condensates. On the molecular level, Tau molecules undergo peptide chain extension during condensation, which promotes interactions between Tau repeat domains and primes condensed Tau for mostly anti-parallel, non-beta-sheet stacking. Over time, however, the same peptide chain extension enables the formation of few beta-structure containing seeds in “aged” Tau condensates. Our data suggest that in cells this Tau condensate maturation process can occur on microtubules or at the nuclear envelope.

## Results

### Loss of molecular diffusion inside and across the interface of Tau/RNA/PEG condensates

To get a more detailed picture of the Tau condensate maturation process, we started by measuring the fluorescent recovery after photobleaching (FRAP) of Tau inside condensates, which can probe how the mobility (∼diffusion) of fluorescently labeled proteins inside condensates changes over time. FRAP after full-condensate bleaching relies on molecules entering the dense condensate phase from the surrounding dilute phase, whereas FRAP after partial condensate bleaching can be achieved via internal redistribution of molecules, without the need of new molecules entering the condensed phase. From FRAP curves, the diffusion of molecules in condensates can be inferred, for example as time to half recovery, t_1/2_ (long t_1/2_ = slow recovery = slow diffusion), and the fraction of mobile molecules is given by the final plateau height.

We performed FRAP on Tau/RNA/PEG condensates (10 μM recombinant human full-length Tau (2N4R isoform) with 1% fluorescently labeled Tau-Alexa488, 0.1 μg/ml polyA RNA, 5% PEG8000) deposited on a microscope slide at different time points (1 h, 4 h, and 24 h) after their formation. When bleaching parts of Tau/RNA/PEG condensates (∼10% of condensate area), both the fluorescent recovery rate and the mobile fraction of Tau molecules gradually decreased with condensate aging (Fig. 1B), indicating the progressive formation of interactions inhibiting the movement of Tau molecules inside condensates. When bleaching entire condensates, we observed increasing recovery times but similar mobile Tau fractions (∼30%) for 15 min, 1 h, and 4 h-old condensates (Fig. 1C). However, after 24 h, few mobile molecules remained and recovery was very slow, indicating almost full loss of Tau’s molecular mobility within 24 h-old condensates.

**Fig. 1.**
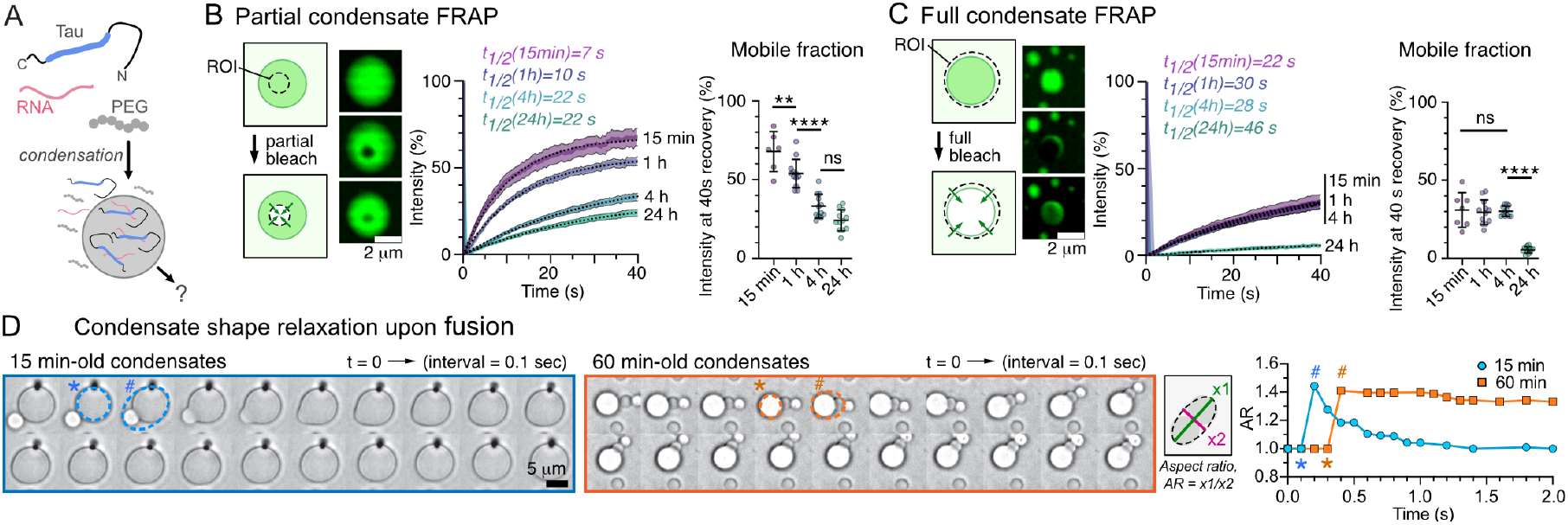
Molecular diffusion in aging Tau/RNA/PEG condensates. **(A)** Schematic of Tau condensation in presence of RNA and PEG**. (B)** FRAP of Tau after partial bleaching (ROI ∼10% of condensate) at different times (15 min, 1 h, 4 h, 24 h) after Tau/RNA/PEG condensate formation. Example images and FRAP curves show recovery of fluorescence over time. Black dotted lines indicate single exponential fits to each condition, t_1/2_ as indicated. Graph on the right shows significance of final recovery after 40 s between different time points. N = 12-13 condensates per condition from 3 experiments. Data shown as mean±SD. One-way ANOVA with Tukey post-test. Scale bar = 2 µm. **(C)** Fluorescence recovery after photo bleaching (FRAP) of Tau after full bleaching (ROI ∼100% of droplet size) at different times (15 min, 1 h, 4 h, 24 h) after Tau/RNA/PEG condensate formation. Example images and recovery curves show recovery of fluorescence over time. Black dotted lines indicate single exponential fits to data, t_1/2_ as indicated. Graph shows significance of final recovery after 40 s. N = 12-13 condensates per condition from 3 experiments. Data shown as mean±SD. One-way ANOVA with Tukey post-test. Scale bar = 2 µm. **(D)** Time lapse of condensate fusion recorded for 15 min and 60 min-old Tau condensates. Condensates were brought in proximity using the laser trap (black trap bead in top panel), while fusion events were unassisted. Recording the aspect ratio (AR = long axis (x1)/short axis (x2) of fitted ellipse) of two fusing condensates shows full recovery of circular shape (AR = 1) for 15 min-old condensates in 2 s. 60 min-old condensates do not fully fuse (AR remains > 1). In image and graph, * = condensate right before, and ^#^ = right after initial fusion. Scale bar = 5 µm.

Time-lapse imaging of condensate fusion events (Fig. 1D) showed that fresh (15 min-old) Tau/RNA/PEG condensates would fuse and re-adopt a circular shape (aspect ratio, AR = 1) within seconds, whereas 1 h-old condensates attach to each other but did not achieve complete fusion and shape relaxation (AR > 1). Together these data confirmed our previous observations of declining Tau mobility in maturing Tau/RNA/PEG condensates (*11*), and, additionally, suggested a complex polymerization process.

### Elasticity dominated, viscoelastic “aging” of Tau condensates

Condensates formed by soluble proteins have been shown to behave like viscoelastic fluids that during maturation increase in viscosity and maintain their elasticity (Maxwell fluid behavior) (*15*, *16*). During condensate aging, intermolecular interactions inside condensates gradually stabilize, which can lead to dynamic arrest of their constituent molecules. This decrease in molecular mobility results in a change in their viscoelastic properties - called condensate “aging”.

To understand the maturation process of Tau/RNA/PEG condensates and their viscoelastic phase transitions, we performed an active micro-rheology assay using a dual trap configuration in an optical tweezer experiment (*17*). We trapped two polystyrene microspheres and brought them into contact with the equatorial plane of condensates formed from Tau and RNA (Tau/RNA/PEG condensates; d ≈ 7-8 μm; 10 μM human Tau, 10 μg/ml polyA RNA, 20 mM HEPES pH7.5, 10 mM NaCl, 5% PEG8000). The two trapped microspheres firmly adhered to the droplet on opposite sides in the equatorial region (Fig. 2A). While one microsphere was oscillated with varying frequencies (actuator bead), the other was kept static and held the droplet in place to measure the transmitted stress (*15–17*) (Fig. 2B). When pushing the actuator bead towards or away from the static bead, the force exerted on the static bead depended on the viscoelastic properties of the probed condensate. For Tau/RNA/PEG condensates, steady displacement of the actuator bead led to steady force responses of the static bead, indicating purely elastic behavior. For comparison, MEC- 2 condensates, known for their liquid-like viscoelastic behavior (*16*), showed progressive dissipation of the force acting on the static bead, indicating liquid-like viscoelastic behavior (Fig. S1A). Sweeping across increasing oscillation frequencies, ranging from 0.1-64 Hz, we calculated the frequency-dependent response of the droplet from the forces acting on the microspheres, which bears information about the stored and dissipated mechanical stress (*15*, *16*). Assuming a linear response (= small deformation) to small forces (10 pN), we estimated the shear modulus by accounting for surface tension from the droplet’s response function ((*15*, *18*); see Methods). This complex modulus comprises a storage component (G’), reflecting elastic energy stored and recovered during deformation, and a loss component (G’’), reflecting energy dissipated through viscous flow. When G’ exceeds G’’, the material behaves predominantly as an elastic solid (*19*). During the first 4 h after condensate formation, the response of Tau/RNA/PEG condensates was dominated by the storage modulus across the 0.1–64 Hz frequency range, indicating a primarily elastic material (Fig. 2C).

**Fig. 2.**
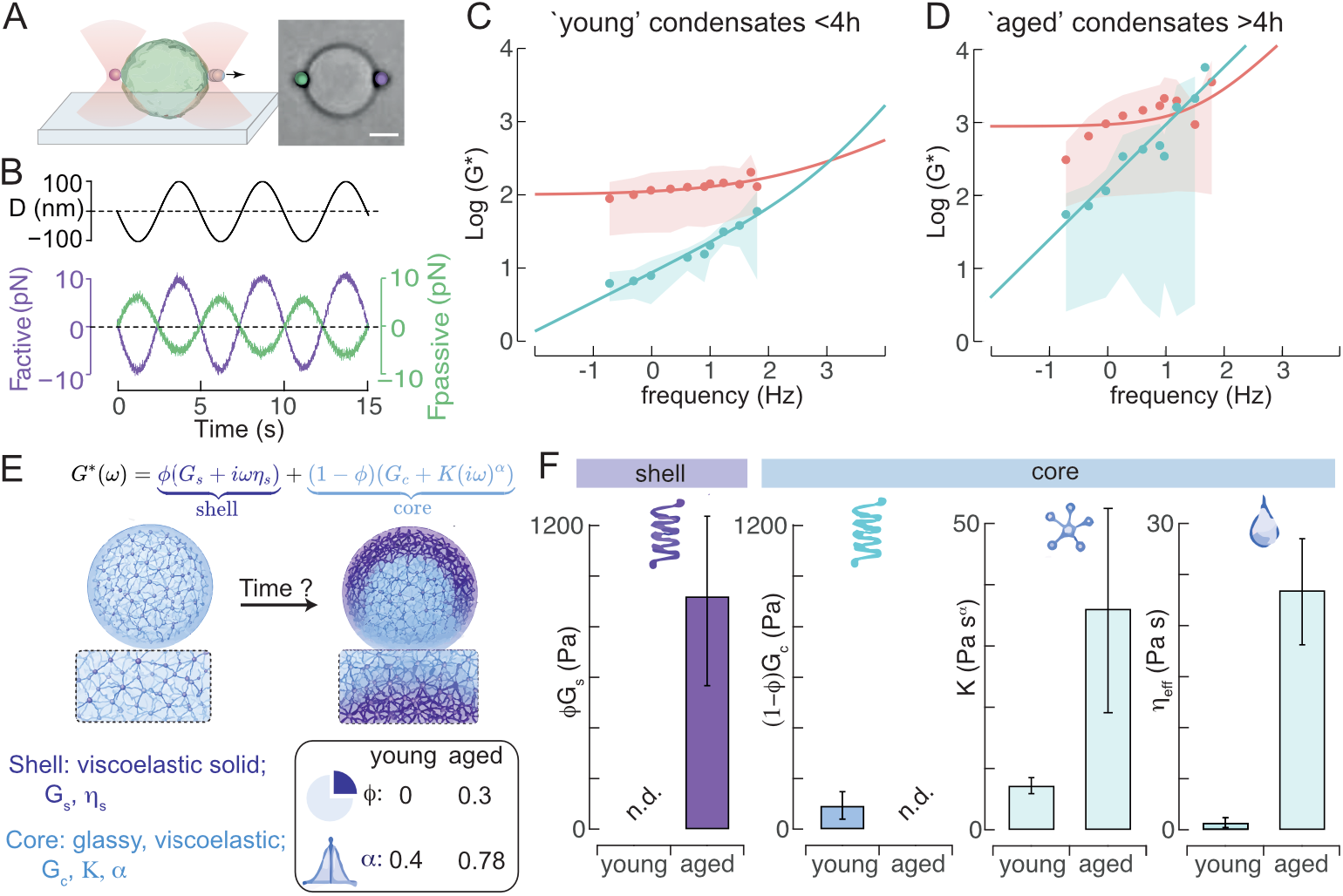
Viscoelasticity of aging Tau/RNA/PEG condensates. **(A)** Schematic of two beads sandwiching an immobilized Tau/RNA/PEG droplet for dual optical trapping experiment. Scale bar = 5 µm. **(B)** Micro-rheology routine showcasing the imposed oscillation (displacement, upper panel) and the corresponding force response on the active (lilac) and the passive (moss) microsphere. Note the small phase difference between active and passive indicates a primarily elastic response. **(C, D)** Frequency-dependent storage (*G*^y^, red) and loss (*G*^yy^, cyan) moduli of naïve (<4 h, N = 26) and mature (>4 h, N = 23) tau condensates, respectively. Solid lines represent fits to the fractional Kelvin–Voigt (fKV) model. Shaded regions indicate the interquartile range of the experimental measurements. The fractional model accurately captures the sublinear power-law dependence of the loss modulus that is not reproduced by the classical Kelvin–Voigt model. **(E)** Schematic of the shell-core composite model. Mature condensates are represented as a mechanically coherent outer shell surrounding a heterogeneous viscoelastic core. The shell is modeled as a Kelvin–Voigt element with elastic modulus *G_s_* and viscosity *η_s_*, whereas the core is described by a fractional Kelvin–Voigt element with elastic modulus *G_c_*, distributed viscoelastic coupling *K*, and fractional exponent *α*. f denotes the fractional contribution of the shell. **(F)** Parameters obtained from the shell-core model. Shown are the effective shell and core elastic contributions, *φG_s_*and (1 − *φ*)*G_c_*, together with the weighted core fractional coupling (1 − *φ*)*K*and the effective viscosity *η*_åçç_evaluated at 1 Hz. Naïve condensates are dominated by a fractional viscoelastic core with a negligible shell contribution (*φ* ≈ 0, *α* ≈ 0.4). During aging, a mechanically significant shell emerges (*φ* ≈ 0.3), accompanied by an increase in shell elasticity, distributed viscoelastic coupling, effective viscosity, and the fractional exponent (*α* ≈ 0.8), consistent with the development of a mechanically coupled cortical layer surrounding a viscoelastic condensate interior. Data shown as mean±SD.

Notably, unlike condensates characterized by a viscoelastic fluid (Maxwell) behavior (*15*), at early time points (<4 h), the naive Tau/RNA/PEG condensates exhibited predominantly viscoelastic solid-like behavior, with the storage modulus (= elastic component) exceeding the loss modulus (= viscous component) across the probed frequency window (*19*). However, the loss modulus increased with frequency much more slowly than the linear dependence predicted for a classical Kelvin–Voigt material. This deviation indicates that stress relaxation is not governed by a single characteristic timescale, but rather by a broad spectrum of relaxation processes better described by fractional viscoelastic models (*20*). This behavior suggests heterogeneous internal dynamics characterized by a broad spectrum of relaxation times, rather than dissipation governed by a single dominant viscous process coupled to an elastic scaffold.

Next, to assess the material aging process, we measured the same samples after >4 h of condensate formation (= “mature” condensates). We found that the complex shear modulus (G* = G’ + iG’’) was significantly increased (Fig. 2D). Again, for all frequencies tested, the droplets behaved as an elastic material with a dominating storage modulus, G’, that increased nearly 10-times over the frequency range tested. This indicated that Tau/RNA/PEG condensates significantly increased in elasticity during maturation. In addition, the loss modulus of aged Tau/RNA/PEG condensates also increased monotonically with probing frequency, but much more quickly than for naïve droplet. These findings suggest that condensate aging is accompanied by the emergence of a more organized internal structure in which tau molecules become increasingly mechanically coupled. Such changes could arise from the formation of a more interconnected molecular network within the condensate and/or from the development of a denser, mechanically reinforced layer at the droplet surface.

To investigate whether the observed rheological evolution could arise from the emergence of spatially heterogeneous mechanics, we next considered a shell-core composite model (Fig. 2E). In this framework, the condensate is represented as a mechanically distinct outer shell surrounding a viscoelastic core,

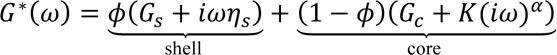

Here, *ϕ* quantifies the relative contribution of the shell to the overall mechanical response, *G_s_* and *η_s_* describe the elastic and viscous properties of the shell, respectively, and *G_c_*, *K*, and *α* characterize the viscoelastic behavior of the condensate interior (see Methods). Specifically, *G_c_* represents the low-frequency elastic plateau modulus of the core and therefore reflects the mechanically persistent fraction of the condensate interior. The parameter *K* determines the magnitude of the fractional viscoelastic response and can be interpreted as a measure of distributed viscoelastic coupling, reflecting the extent to which molecular rearrangements are mechanically linked across the condensate. Finally, the fractional exponent *α* describes the distribution of relaxation timescales within the core (*20*). Low values of *α* indicate highly heterogeneous, glassy-like dynamics characterized by a broad spectrum of relaxation times, whereas larger values indicate increasingly coordinated and network-like mechanical behavior while remaining distinct from classical single-timescale viscoelasticity.

Consistent with the absence of a cortical shell at early time points, fitting converged to a negligible shell contribution (ϕ≈0), effectively reducing the model to a homogeneous fractional Kelvin–Voigt material. In contrast, aged condensates exhibited a finite shell contribution (ϕ≈0.31), indicating that approximately one-third of the measured mechanical response could be attributed to a mechanically distinct outer layer.

Notably, the shell elasticity emerged (*G_s_*≈880 Pa), whereas the shell viscosity converged toward zero, suggesting that the shell behaves predominantly as an elastic scaffold rather than as a conventional viscoelastic layer (Fig. 2F). Within the core, aging was accompanied by increases in both *K (K_early_* = 7 ± 2 to *K_late_ =* 36± 13; mean±SD) and *α (α_early_* = 0.39 ± 0.05 to *α_late_ =* 0.78± 0.09; mean±SD), indicating a transition from highly heterogeneous dynamics toward increasingly coordinated viscoelastic deformation. Together with the emergence of a finite shell contribution, these results suggested that Tau condensates undergo mechanical maturation through both internal network organization and cortical reinforcement, resulting in a spatially heterogeneous shell-core architecture.

### Internal restructuring of Tau during condensate maturation

To elucidate the molecular restructuring leading to the increase in Tau condensate elasticity and “shell” formation during maturation, we applied optical diffraction tomography (ODT), which determines the mass density distribution at microscopic length scales (*21*, *22*). We collected refractive index (RI) tomograms of Tau/RNA/PEG condensates at 1 h, 4 h, and 24 h after preparation (Fig. 3A). 1 h-old Tau/RNA/PEG condensates showed a homogeneous mass density distribution in their interior (Fig. 3B; equatorial plane of condensates), with a total Tau protein and RNA concentration of about ∼220 mg/ml in the center of the condensates (average radial plots of condensates; Fig. 3C,D). After 4 h, the concentration of material inside condensates increased to ∼260 mg/ml whereby the condensate interior mostly maintained a homogenous distribution of mass density. 24 h-old condensates on average had a larger diameter, a substantially lower central mass density of ∼175 mg/ml, a higher density of ∼200 mg/ml at the outer border of the condensates, and showed an inhomogeneous material distribution. In profile plots of individual 0.5 μm-thick equatorial plane projections, dense nodes alternated with regions of low density in 24 h-old condensates (Fig. 3C). This inhomogeneous material distribution was also apparent from the large standard deviation of the average radial densities at 24 h (Fig. 3D). Equatorial plane ODT images of different 1 h, 4 h, and 24 h-old condensates showed that the mass density distribution inside condensates mostly homogenously increased between 1 h and 4 h, but was highly variable in 24 h-old condensates, with pronounced density irregularities in most condensates, characterized by nodes of high density and void spaces lacking material, reminiscent of porous particles (Fig. 3E). Furthermore, we observed apparent internal fibrillar networks and a shell-like compaction of material at the condensate surface, supporting the data and fit model derived for our viscoelasticity measurements. For a few condensates, the formation of fibrillar structures could be observed at the interface of condensed and dilute phase, protruding into the dilute phase (Fig. 3E).

**Fig. 3.**
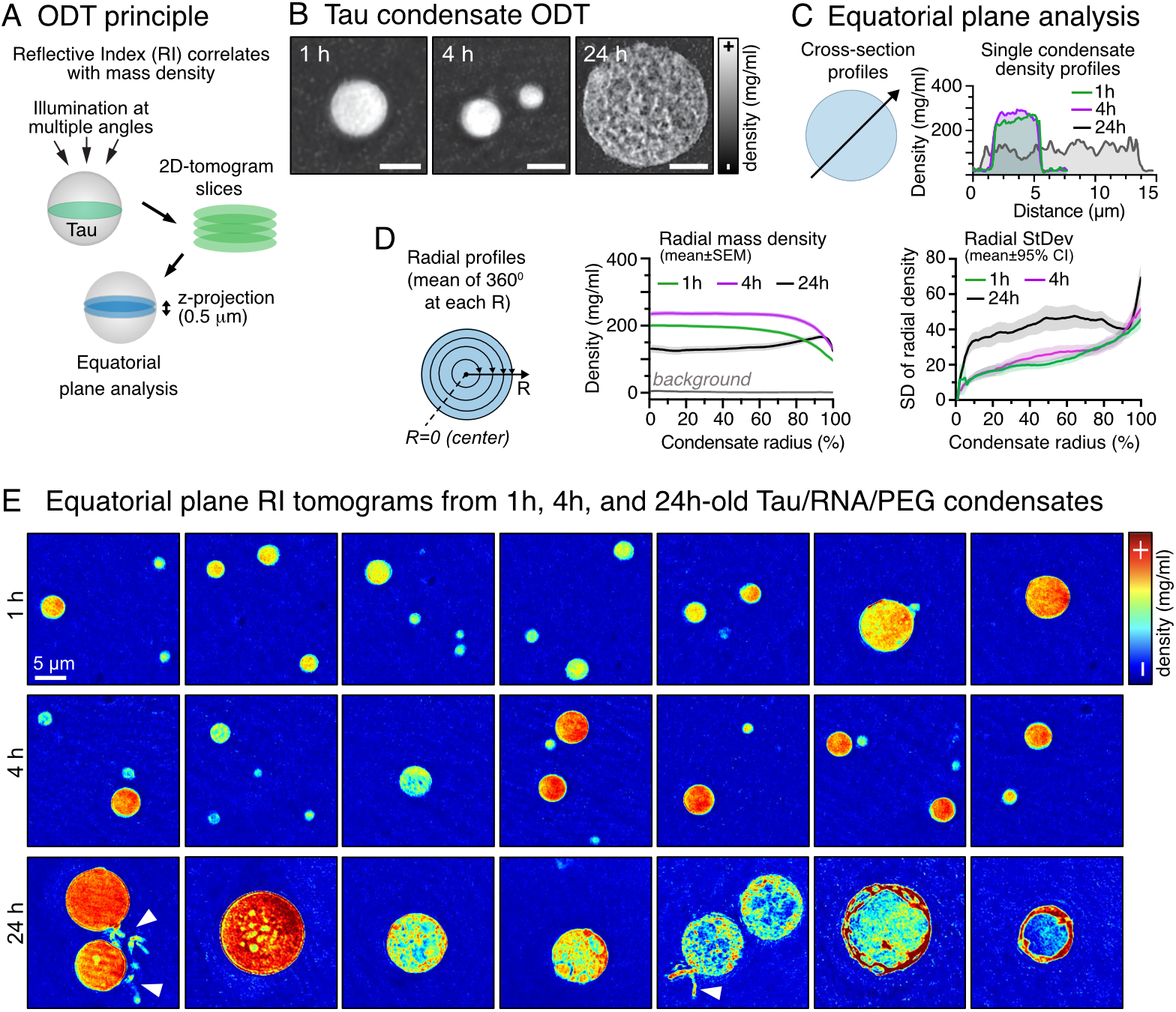
Molecular restructuring during the maturation of Tau/RNA/PEG condensates. **(A)** ODT using tomograms of refractive index (RI) maps to reconstruct the density distribution in the equatorial plane of Tau/RNA/PEG condensates. **(B)** Example ODT and fluorescence images of the equatorial plane of 1 h, 4 h, and 24 h-old YFP-Tau/RNA/PEG condensates. Scale bars = 5 μm. **(C)** Line profiles plots through equatorial plane of individual example condensates. **(D)** Average radial mass density profiles of Tau/RNA/PEG droplets at 1 h, 4 h, and 24 h. Mass density (mg/ml) is plotted from the center (R = 0%) to the border (R = 100%) of condensates. Radial variance plot shows the average variance in density measured for each condensate at a given radius. N = 33-56 condensates per condition, radial mass density data shown as mean±SEM, radial standard deviation (StDev) data shown as mean±95% confidence interval. **(E)** Gallery of equatorial plane density maps of different example 1 h, 4 h, and 24 h-old Tau/RNA/PEG condensates. White arrow indicates fibrillar structures protruding from the condensate surface. Scale bar = 5 μm.

To assess whether the formation of structure inside Tau/RNA/PEG condensates depended on the presence RNA, we performed ODT experiments on aging Tau/PEG condensates lacking RNA (Fig. S2A). At 1 h, Tau/PEG condensates, which were substantially smaller than Tau/RNA/PEG condensates, displayed a homogeneous mass density distribution with an average density of ∼100 mg/ml in their center (Fig. S2B,C,D), which remained similar in 4 h-old Tau/PEG condensates. Some 4 h-old condensates had increased densities of ∼200 mg/ml. At 24 h, Tau/PEG condensates displayed markedly non-circular, irregular shapes and a higher central mass density of ∼300 mg/ml. Unlike Tau/PEG/RNA condensates, Tau/PEG condensates did not develop shell-like compaction at the condensate surface; instead, densification occurred preferentially at their core. Furthermore, many non-spherical 24 h-old condensates exhibited two distinct density maxima within their interior (arrows in Fig. S2B,D), reminiscent of two incompletely fused condensates.

Together, these data suggest that the presence of RNA promotes the inhomogeneous restructuring of Tau inside condensates, leading to the formation of fibrillar structures and the formation of a core-shell arrangement. In the absence of RNA, Tau polymerization seems to be slower and homogenous inside condensates.

### Progressive molecular compaction and Tau-Tau interactions in aging Tau condensates

To test to which extend the restructuring inside Tau/RNA/PEG condensates involved Tau molecules, we measured CFP-tagged Tau interactions by fluorescent lifetime imaging (FLIM). Conceptually, the lifetime of CFP should decrease with increasing CFP-Tau density in condensates, and even further upon Tau aggregation due to fluorescence “self-quenching” effects (*11*, *23*) (Fig. S3A). Different Tau density domains inside condensates, as suggested by ODT, would appear as inhomogeneities in CFP lifetimes. In preparations of CFP-Tau/RNA/PEG condensates, we found that the CFP lifetime (LT) was significantly reduced inside (dense phase; LT_dense_) compared to the outside (dilute phase; LT_dilute_) of condensates (Figs. S3B,C), which is expected because of the high density of CFP-Tau molecules in the dense compared to the dilute phase.

In addition, it appeared that different molecular CFP LTs, and therefore CFP-Tau densities, coexisted inside condensates, apparent from different LT components (LT_dense-1_ (yellow) and LT_dense-2_ (cyan)) assignable to the interior of CFP-Tau/RNA/PEG condensates from FLIM phasor plots (see methods). 24 h-old condensates appeared to have more or larger LT_dense-2_ clusters, and both LT_dense-1_ and LT_dense-2_ inside CFP-Tau/RNA/PEG declined over time (Fig. S3C), suggesting that the fraction of denser condensate domains as well as the Tau density in all condensate domains increased over time, reminiscent of the progressive, inhomogeneous densification observed by ODT.

To test whether increasing Tau-Tau interactions were responsible for the decay of CFP LTs during condensate maturation, we performed FLIM experiments on Tau/RNA/PEG condensates formed with a 1:1 mix of CFP-Tau and YFP-Tau (Fig. S3D). CFP LTs inside CFP/YFP-Tau condensates were generally lower because of CFP/YFP FRET and showed a CFP LT decline mostly between 4h and 24h, which could be related of the molecular restructuring and shell formation occurring in this time window (compare ODT and rheology data).

Interestingly, we also observed a decrease in CFP lifetime in the light phase, LT_dilute_, for both CFP-Tau and CFP/YFP-Tau condensates over time, which may result from the formation of Tau dimers (*24*) and small clusters (radius ∼100-200 nm) (*11*). Overall, CFP-Tau LT imaging supported that the density inhomogeneities detected during condensate maturation by ODT were due to rearrangement and polymerization of Tau molecule inside condensates; high-density regions observed in 24 h-old condensates therefore likely contained densely packed Tau.

### Loss of RNA during Tau/RNA condensate maturation

Radial density profiles from ODT measurements indicated a modest but significant increase in condensate density from 1 h to 4 h, and a decreased density of 24 h-old compared to 1 h- and 4 h-old condensates (Figs. 3C,D). To better understand the cause of these apparent changes in density, we reconstructed whole condensate from ODT tomograms (Fig. 4A). This showed that, from 1 h to 4 h, both total condensate volume and dry mass increased non-significantly. However, as volume doubled but mass tripled, this led to an overall significant increase in average condensates density (= mass / volume; p<0.001) (Fig. 4B; Fig. S3). For 24 h-old condensates, total condensate volume and dry mass further increased compared to 4h-old condensates, while condensate density decreased. These observations suggested that the Tau concentration inside condensates first increased and later decreased – concomitant with the observed internal restructuring in Tau/RNA/PEG condensates. Graphing condensate volume versus density for individual condensates (Fig. 4C) further confirmed the decrease in density at continued condensate growth in 24 h-old condensates. This observation indicated the absence of recruitment or even a net loss of molecules/”material” – Tau and/or RNA molecules – after condensate growth has been (largely) completed, or the influx of water during condensate growth. Since this observation coincided with reduced Tau FRAP (Fig. 1) and internal restructuring, fibrilization and shell formation of Tau molecules inside condensates (Fig. 2 and 3), it marked a shift in the Tau/RNA/PEG condensate maturation process from common condensate growth and densification (i.e., by Oswald ripening (*25*) and homogeneous polymerization) towards pronounced macromolecular restructuring and inhomogeneous fibrillar polymerization.

**Fig. 4.**
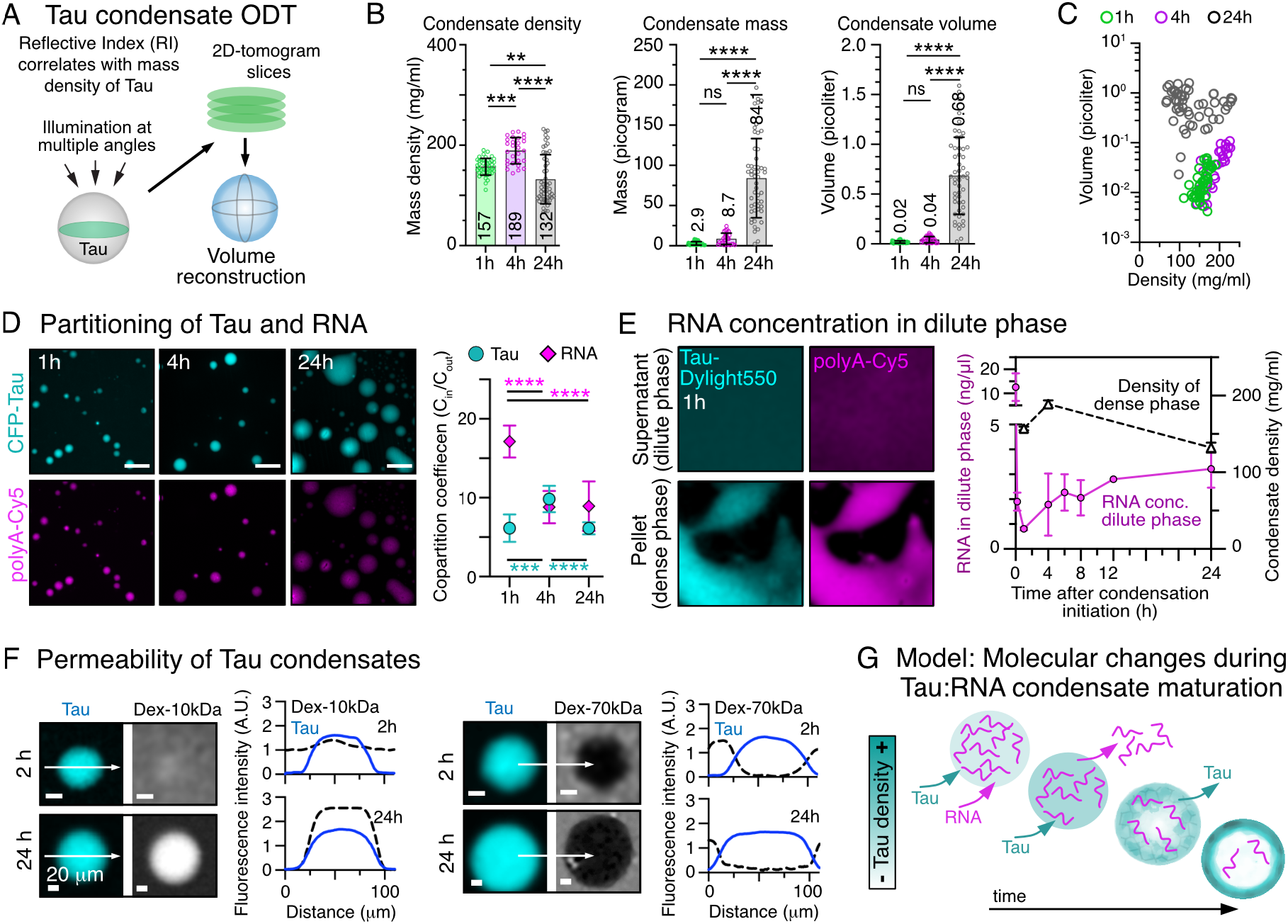
Tau restructuring enables loss of RNA from. Tau/RNA/PEG **condensates. (A)** ODT using tomograms of refractive index (RI) maps to determine volume, density, and dry mass of whole Tau/RNA/PEG condensates. **(B)** Whole condensates density (mg/ml), volume (picoliter), and total mass (picogram) of Tau/RNA/PEG condensates at 1, 4, and 24 h after formation. Data shown as mean±SD, N = 28-54 condensates per time point. Mean values are indicated. One-way ANOVA with Tukey post-test. **(C)** Scatter plot of condensate volume vs. density. N = 28-54 condensates per condition. **(D)** Confocal images of fluorescently labeled Tau (cyan) and RNA (pink) of Tau/RNA/PEG condensates at 1, 4, and 24 h after formation. Scale bar = 5 µm. Quantification of Tau and RNA fluorescent co- partition coefficients ([c_in_]/[c_out_]) indicate a decrease in RNA concentration in 24 h-old condensates. Data shown as mean±SD, N = 4 images per condition. **(E)** Images of Tau (cyan) and RNA (pink) in the dense phase pelleted by centrifugation, and in the remaining light phase. Graph: RNA concentration (pink) measured over time in the light phase as absorption at λ = 260 nm. Data shown as mean mean±SD, N = 3 independent experiments. Average condensate density (black; ODT data from Fig. 3B) are shown for comparison. **(F)** Fluorescently labeled dextran-10kDa and dextran-70kDa was added to 1 h- and 24 h-old Tau/RNA/PEG condensates. Cross-sectional line plots (along white arrows) show differential exclusion and enrichment of the dextrans inside condensates. **(G)** Model: The initial phase of Tau/RNA/PEG condensate maturation is characterized by a further enrichment of Tau and a rapid depletion of RNA. This phase is followed by restructuring and polymerization of Tau molecules inside condensates, which leads to the formation of spheres with a “shell-like” Tau accumulation that remains permeable for RNA and small molecules (∼10 kDa), whereas larger molecules are excluded.

To determine, how the molecular restructuring occurring inside Tau/RNA/PEG condensates between 4 h and 24 h would impact their molecular composition, we determined, we determined the enrichment of fluorescently labeled Tau and RNA in aging Tau/RNA/PEG condensates as co-partitioning coefficient ([fluorescence. intensity_in_]/[fluorescence. intensity_out_] ∼ [c_in_]/[c_out_]). We found a significant decrease of RNA enrichment in condensates from 1 h to 4 h, which then plateaued until 24 h (Fig. 4D). We further confirmed this observation by monitoring the RNA concentration in the dilute phase (absorption at 260 nm) after pelleting the condensates by centrifugation at different time points (Fig. 4E). In contrast to RNA, the enrichment of Tau in condensates increased from 1 h to 4 h and then decreased from 4 h to 24 h (Fig. 4D).

Together, these observations indicate that early phases (first 4 h) of condensate maturation are characterized by a strong loss of RNA and an increase in Tau, followed by a loss of Tau thereafter. These observations suggest that Tau/RNA/PEG condensates must remain able to exchange molecules with the surrounding dilute phase in the first 4 h, which may be abolished at prolonged maturation. The diffusion barrier that we suspected to form around Tau/RNA/PEG condensates within the first hours (compare FRAP data), thus, maybe more permeable for RNA than for Tau. The restructuring of Tau inside condensates during further maturation, i.e., in 24 h-old condensates (compare ODT and optical tweezer data; Fig. 2 and 3), may lead to condensate porosity allowing the exchange of soluble non-polymerized Tau molecules with the surrounding light phase. Accordingly, we observed that the partitioning of fluorescently labeled low molecular weight dextran (MW = 10 kDa; R_H_ ∼ 2 nm; (*26*)) was restricted in 1 h-old condensates but permitted in 24 h-old condensates (Fig. 4F). High molecular weight dextran (MW = 70 kDa; R_H_ ∼ 7 nm; (*26*)) was excluded from both 1 h and 24 h-old condensates. This suggests that aged Tau condensates, despite adopting a porous character, maintained a diffusion barrier for large molecules, whereas Tau, RNA, and smaller molecules can be exchanged with the interior (Fig. 4G).

### Tau molecular extension and anti-parallel organization in young condensates

To understand the molecular interactions underlying Tau condensation and condensate maturation, we performed chemical cross-linking mass spectrometry (XL-MS) coupled with ^15^N-isotopic labeling (*27*, *28*) on 1h-old condensates. (Fig. 5A, Figs. S3A, B). To distinguish intra- from inter-links, condensates were prepared using a 1:1 mix of ^14^N and ^15^N-labeled recombinant Tau, followed by cross-linking under varying conditions. Cross- linked peptides for which only pure ^14^N^14^N or ^15^N^15^N envelopes were detected, with no detectable hybrid ^14^N^15^N / ^15^N^14^N signal, were classified as predominantly intramolecular under our detection threshold. (Fig. 5B).

**Fig. 5.**
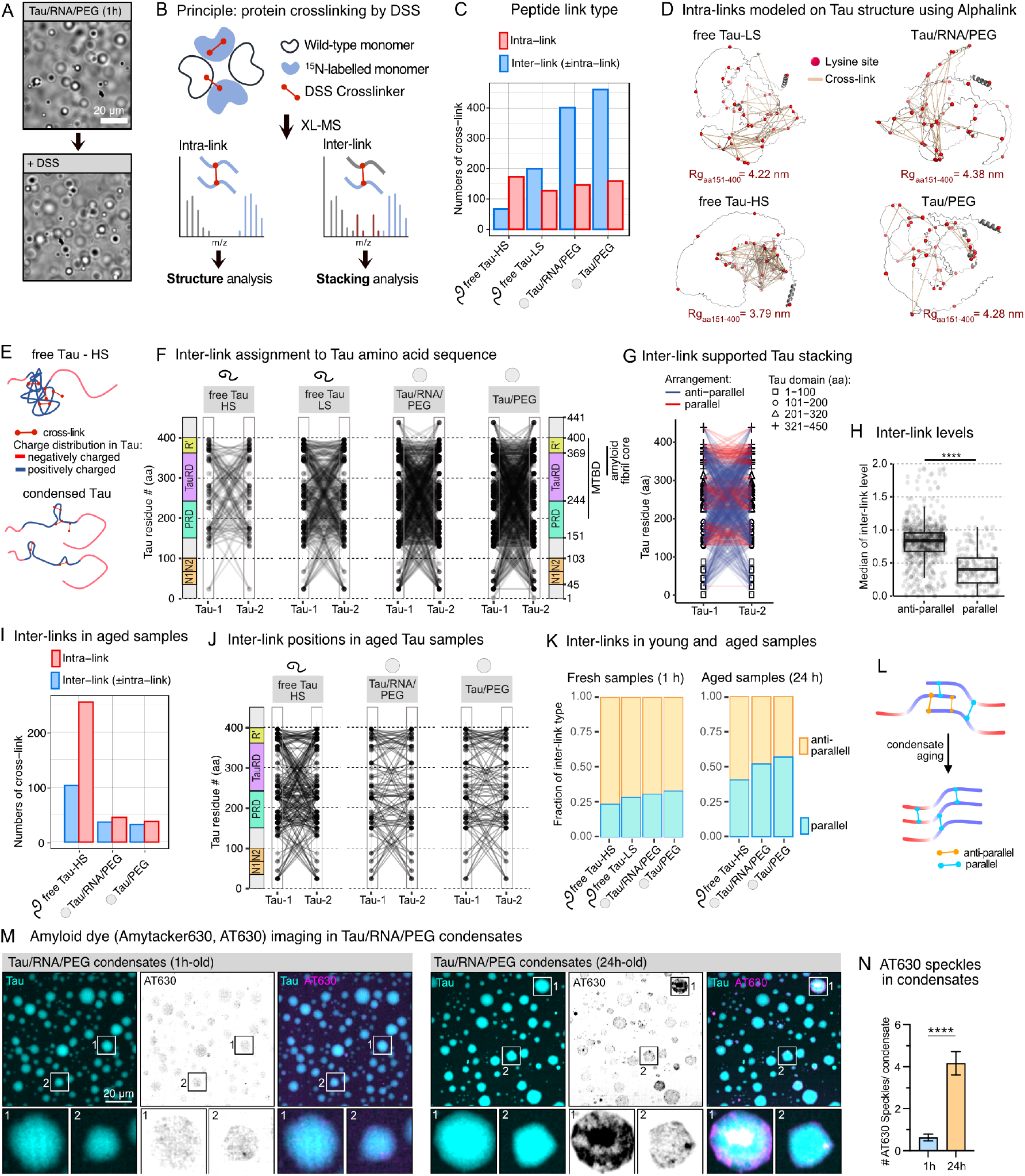
Crosslinking mass spectrometry of young and aged Tau condensates. **(A)** Microscopy of Tau/RNA/PEG condensates before and after DSS crosslinking. Scale bar = 20 μm. **(B)** Principle of intra- and inter-link detection using XL-MS coupled with equally-mixed ^14^N and ^15^N Tau. **(C)** Number of inter- and intra-links in soluble Tau in high salt (HS, 300 mM NaCl) and low salt (LS, 5 mM NaCl) buffer, and in Tau condensed with PEG and RNA (Tau/RNA/PEG) or with only PEG (Tau). **(D)** Tau structure with constraint intra-links using AlphaLink. Position of lysine residues (red dots) in detected cross-links (yellow lines) are indicated. **(E)** Model of Tau conformational extension during condensation based on intra-link analysis. **(F)** Position of inter-links between two Tau molecules, mapped onto the Tau sequence. **(G)** Classification of inter-links between same Tau domains (homotypic domain link = parallel, red) and different Tau domains (heterotypic domain link = anti- parallel, blue) in different Tau molecules. Assignment based on Tau regions of 100 aa. **(H)** Comparison of inter-link level in parallel vs. anti-parallel arrangement using Wilcoxon’s test (adjusted p-value < 0.001). **(I)** Peptide links in aged (24 h-old) soluble Tau and Tau condensates. **(J)** Inter-link mapping in aged Tau and Tau condensate samples. **(K)** Fractions of parallel vs. anti-parallel inter-links in fresh and aged Tau samples. **(L)** Illustration of possible Tau arrangements having mixed anti-parallel and parallel (top) or a predominantly parallel (bottom) stacking. **(M)** Microscopy of Amytracker 630 (AT630) in young (1 h-old) and aged (24 h-old) condensates formed with CFP-Tau. Inserts show selected condensates having larger AT630+ Tau structures in their interior and at their surface at higher magnification. Scale bar = 20 μm. **(N)** Quantification of AT630-positive, bright speckles in 1h- and 24h-old Tau/RNA/PEG condensates. Data shown as mean±SEM, N ∼ 30 condensates per condition. Unpaired Student t-test.

We compared the relative probabilities of inter- and intramolecular Tau cross-links across four conditions: Tau/RNA/PEG/PEG condensates (10 μM Tau, 5 μg/ml RNA, 5% PEG, pH7.4, 5 mM NaCl), Tau/PEG condensates without RNA (10 μM Tau, 5% PEG, pH7.4, 5 mM NaCl), soluble Tau in the same low salt buffer (10 μM Tau, pH7.4, 5 mM NaCl; Tau-LS), and soluble Tau in high salt (10 μM Tau, pH7.4, 300 mM NaCl; Tau-HS) to prevent intramolecular electrostatic interactions and cluster formation. Using MS1 isotope-envelope analysis, we classified cross-links with detectable hybrid species as containing intermolecular contributions, whereas cross-links lacking detectable hybrid envelopes were assigned as predominantly intramolecular (COMPASS analysis, (*27*)). Our analysis revealed that Tau condensation, both with and without RNA, substantially increased the number of inter-molecular links, but not intra-molecular links, relative to soluble Tau (Fig. 5C, Fig. S4C). Conversely, higher ionic strength (300 mM NaCl) decreased inter-links in soluble Tau while increasing intra-links.

To further examine the Tau conformational states represented by the intramolecular cross- links, we modeled these intra-links onto Tau structure predictions using AlphaLink 1.0.0 (*29*). Compared with AlphaFold, AlphaLink substantially reduced the Cα–Cα distances of mapped intra-links while preserving the overall low-confidence profile expected for intrinsically disordered Tau (Fig. S4D), supporting the use of XL-informed modeling to evaluate condition-dependent Tau conformations. AlphaLink modeling of full-length Tau, the C-terminal part of Tau (TauC, aa 151-441) and the Tau repeat domain (TauRD, aa 244- 372) revealed a pronounced compaction (∼ modeled radius, Rg) of soluble TauC and TauRD under high-salt conditions (Fig. 5D, E; Fig. S4E). Conversely, condensation seemed to promote more extended TauRD conformations in Tau/PEG condensates (Fig. 5E, Fig. S4E) and induced significantly larger average Cα–Cα cross-link distances (∼50 Å) within Tau/RNA/PEG condensates exhibited than those in soluble Tau and Tau/PEG condensates (Fig. S4F). Notably, using AlphaLink to model and constrain Tau’s structure, the average Cα-Cα distances (∼50 Å) detected in Tau/RNA/PEG condensates violated the maximum distance between two lysine side chain cross-linked by DSS (∼24 Å). These apparent restraint violations indicate that the condensate-derived cross-linking pattern cannot be represented by a single static Tau conformation. Instead, they are consistent with the highly dynamic and heterogeneous ensemble behavior of Tau in condensates, where transiently populated conformations and intermolecular contacts contribute to the observed cross- linking pattern.

To investigate inter-molecular Tau interactions within condensates, we mapped the detected inter-links onto two juxtaposed Tau peptide chains (Fig. 5F). As anticipated, the lysine-rich proline-rich domain (PRD), repeat domain (TauRD), and R’ (amino acids (aa) 370-400) accounted for the majority of inter-links across all conditions. Notably, the C-terminal region (> 400 aa) formed cross-links in condensates but not in free Tau. Although condensation substantially increased the total number of inter-links compared to free Tau, the specific residue positions involved in inter molecular contacts showed few systematic differences. The pattern of both homotypic inter-links within the same domains and heterotypic inter-links spanning distinct, distant domains could not be explained by a single molecular arrangement, implying the coexistence of at least two distinct stacking modes. A parallel, linearized Tau arrangement can largely account for the high frequency of homotypic inter-links within the PRD, TauRD, and R′ domains (Fig. 5G). However, this arrangement would impose geometric constraints that are incompatible with the simultaneously observed long-range heterotypic inter-links, which require antiparallel and/or more compact configurations to bring distant regions into proximity. Thus, the coexistence of extensive homotypic and heterotypic inter-links across the dataset suggests that Tau molecules do not adopt one uniform stacking geometry, but instead populate multiple stacking configurations within condensates.

To further investigate the structural arrangements of Tau within condensates, we quantified and compared inter-link levels corresponding to parallel versus anti-parallel alignments. The inter-link signal for the anti-parallel arrangement was significantly higher than that for the parallel arrangement (Wilcoxon test, p.adj < 0.001). Furthermore, the number of unique residue pairs forming anti-parallel inter-links exceeded those forming parallel inter-links. Notably, the proportion of parallel arrangement inter-links showed a gradual increase during the early stages of condensation (Fig. 5F, H).

### Evolution of parallel Tau stacking during condensate maturation

To investigate the molecular rearrangements associated with condensate maturation, we extended our XL-MS analysis to 24 h-old Tau condensates (Fig. S4G). The number of identifiable cross-links in aged condensates was substantially lower (< 50) than in 1 h-old condensates (> 100) (Fig. 5I), a phenomenon likely resulting from over-crosslinking and the formation of a higher-order cross-linked network resistant to enzymatic digestion.

Consistent with this interpretation, the precursor ion mass distribution in aged condensates shifted toward 4000–5000 Da, compared to 2500–3500 Da for free Tau (Fig. S4H). Furthermore, while 1-h old condensates predominantly yielded four isotopic peak envelopes (¹⁴N¹⁴N, ¹⁴N¹⁵N, ¹⁵N¹⁴N, ¹⁵N¹⁵N) expected from a 1:1 ¹⁴N/¹⁵N mixture, precursors from 24 h-old condensates frequently exhibited more than eight envelopes (e.g., ¹⁴N¹⁴N¹⁴N, ¹⁴N¹⁴N¹⁵N, ¹⁴N¹⁵N¹⁴N), indicative of higher order cross linked species (Fig. S4I). Similar patterns were observed in recombinant amyloid-like Tau fibrils, suggesting that Tau molecules in mature condensates form densely and stably packed arrangements analogous to fibrils, which hinder protease digestion. Despite the lower overall identification rate, the residue pairs we could extract revealed a clear trend in the stacking pattern: the proportion of parallel Tau arrangements in aged Tau/RNA/PEG and Tau/PEG condensates increased to >50%, compared to <30% in young condensates (Fig. 5J, K). Collectively, these data indicate that during condensate maturation, Tau molecules become progressively more organized, adopting a predominantly parallel arrangement (Fig. 5L).

The core of pathological Tau amyloid fibrils is built from parallel, in-register TauRDs (*30*). The parallel Tau arrangement evolving during condensate maturation could favor amyloid-like Tau stacking and the formation of beta-sheet-rich, seeding competent Tau species. Indeed, staining Tau/RNA/PEG condensates with a dye showing bright fluorescence upon binding to amyloid-containing protein assemblies (Amytracker630, AT630) indicated bright fluorescent speckles inside and on the surface of aged (24 h-old) but not young (1 h- old) condensates (Fig. 5M,N). Only few aged Tau/RNA/PEG condensates showed AT630+ polymerized Tau structures seen by ODT. In aged Tau/PEG condensates, substantially less AT630+ foci could be detected (Fig. S4J). This indicated that Tau/RNA/PEG condensates were especially efficient in catalyzing the formation of oligomeric, beta-structure containing Tau seeds, and that the pronounced structural organizing of Tau in aged Tau/RNA/PEG condensates may only in part resemble amyloid-like Tau structures.

### Evolution of cellular Tau assemblies seeded by aged Tau condensates

Next, we confirmed that 24 h-old Tau/RNA/PEG condensates, which showed amyloid-like Tau foci, could induce cellular Tau aggregation. For this, we transfected aged Tau condensates into Tau biosensor cells – HEK293 cells stably expressing TauRD with the P301S FTD mutation and C-terminally fused to CFP (TauRD^P301S^-CFP; (*31*) ; Fig. 6A) – and monitored the formation of Tau aggregates. Intracellular Tau aggregates formed upon cytosolic delivery of aged, seeding-competent Tau/RNA/PEG condensates and to a lesser extend upon delivery of aged Tau/PEG condensates, often close to the nucleus, and in some cells also inside the nucleus (Fig. 6B,C); non-seeded cells or cells exposed to seeding-incompetent young Tau/RNA/PEG or Tau/PEG condensates did not develop aggregates (Fig. S5; (*11*)). Confocal z-stack time-lapse imaging of the same cells over the course of 15 h (starting 6 h post condensate transfection) revealed the time course of aged condensates- induced Tau aggregation (Fig. 6D,E): first small Tau foci appeared dispersed in the cytosol (CLUS), followed by an accumulation of Tau foci at and around the nucleus (NE), and, later, the formation of larger cytosolic Tau aggregates (CYT) adjacent to the nucleus. The small Tau foci in the cytosol and at the nucleus therefore represent the earliest microscopically visible, condensate-seeded Tau assemblies. High resolution microscopy (STED) revealed that cytoplasmic TauRD “foci” in part resemble thin Tau fibrils that are attached or adjacent to microtubules (MTs) – at least in later stages of aggregation when cytoplasmic aggregates had formed as well –, and showed the localization of Tau foci on the cytoplasmic side of the nuclear envelope (Fig. 6F).

**Fig. 6.**
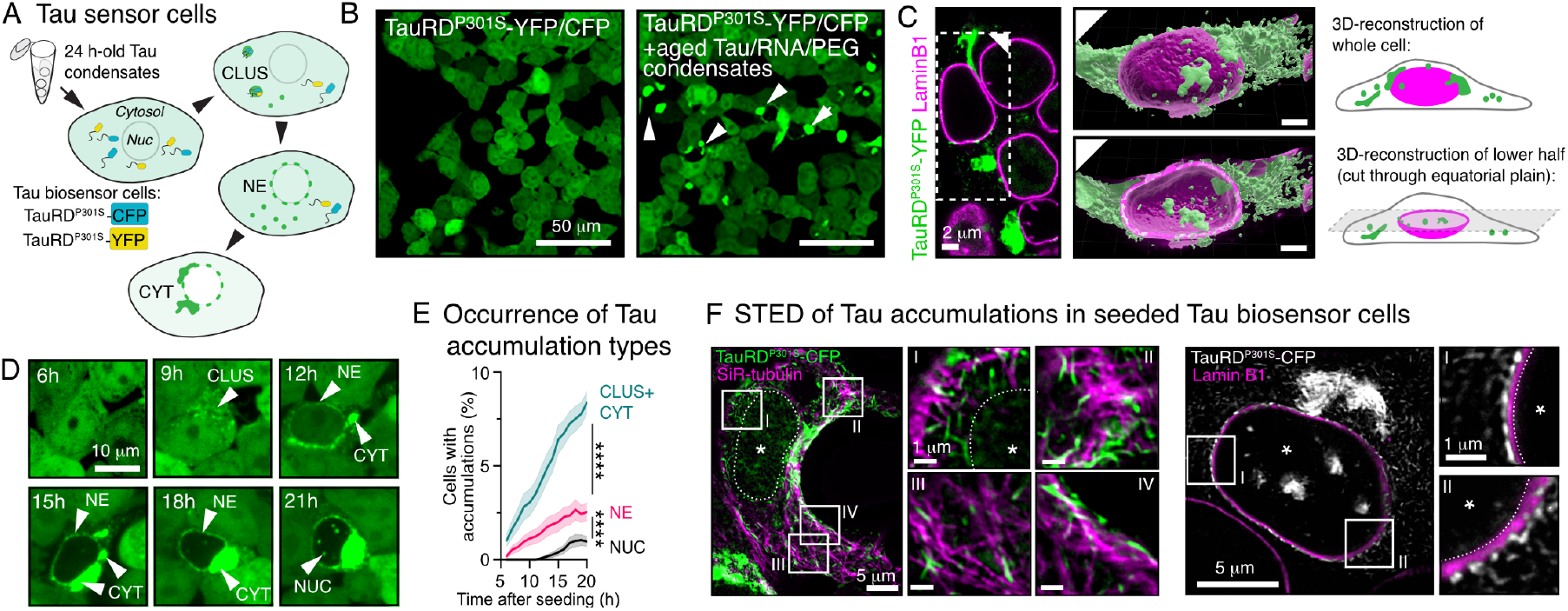
Seeding of cellular Tau aggregation by aged Tau/RNA/PEG condensates. **(A)** Schematic of seeding Tau aggregation in Tau biosensor (HEK293 expressing TauRD^P301S^- CFP) cells by aged Tau/RNA/PEG condensates. **(B)** Example images of Tau biosensor cells seeded, or not, with 24 h-old Tau/RNA/PEG condensates. Scale bars = 50 μm. **(C)** High-resolution imaging and 3D-reconstruction of TauRD^P301S^-CFP in condensate seeded Tau biosensor cells, with counterstaining of the nuclear envelope by Lamin B1 immunostaining shows subcellular positioning of seeded Tau species: Many small Tau foci form in the cytosol and some at the nuclear envelope, larger cytoplasmic Tau aggregates are positioned close to the nucleus, and some Tau clusters also form in the nucleus. Scale bars = 2 μm. **(D)** Confocal time course imaging of Tau biosensor cells upon seeding with 24 h-old Tau/RNA/PEG condensates. Images show sequential formation of Tau accumulation in the same cell: first, cytosolic Tau foci (CLUS) form, followed by Tau foci at the nuclear envelope (NE), larger cytoplasmic Tau aggregates (CYT) close to the nucleus, and, finally, intranuclear circular Tau aggregates (NUC) can be observed. **(E)** Quantification Tau accumulation types from time course imaging experiments. For analysis, cytoplasmic CLUS and CYT were combined. n=21 analyzed time course series (z-stack), data shown as mean±SEM, one-way ANOVA with Tukey post-test for percentage at 21 h for each accumulation class. **(F)** STED microcopy of seeded Tau biosensor cells, counter stained with SiR-tubulin (left panel) or immunostained for Lamin B1 (right panel), showing different Tau accumulation types. Zoom-ins show elongated cytosolic Tau structures adjacent to microtubules (left) and Tau foci at the outer nuclear envelope (right). Position of nuclei are indicated by white stars, inner nuclear envelope-nucleoplasm border is indicated by white, dashed lines. Scale bars = 5 μm in overview and 1 μm in zoom-ins.

Notably, later in the aggregation time course, the formation of intranuclear, spherical Tau aggregates (NUC) occurred in Tau biosensor cells with cytosolic Tau foci and larger aggregates. They may have arisen after cytosolic Tau has compromised the integrity of the nuclear envelope (*11*, *32*) or after nuclear envelope re-assembly after mitosis, both giving seeding-competent Tau oligomers access to nucleate TauRD^P301S^-CFP aggregation in the nucleoplasm. Such intranuclear Tau aggregates are not observed in cells and neurons expressing full-length Tau, where Tau’s access to the nucleoplasm is very limited (*33*); they may therefore be interpreted as an artifact of the Tau biosensor cell line.

### Differential Tau densities of seeded cellular Tau assemblies

To gain insight into the molecular assembly state of Tau foci at the nuclear envelope, we performed FLIM on Tau biosensor cells. FLIM of Tau^P301S^-CFP in seeded Tau biosensor cells suggested that cytosolic and nuclear envelope-associated Tau foci had average CFP lifetimes that were significantly shorter than soluble cytosolic Tau but longer than those observed in cytosolic Tau aggregates (Fig. 7A,B,C). This suggested that Tau foci in the cytosol (CLUS) and at the nuclear envelope (NE) had a more compact structure (= more quenching) than soluble cytosolic Tau. However, these foci appeared to be less compact than larger cytosolic aggregates (CYT), which displayed even lower CFP lifetimes, indicative of amyloid-like structures (AMY). To test whether the reduced fluorescence lifetime of Tau foci and aggregates originated from enhanced interactions of TauRD-TauRD interactions, we performed FLIM in Tau biosensor cells expressing both CFP- and YFP- tagged TauRD^P301S^. In these cells, Tau foci and aggregates had even lower CFP-lifetimes, which can be attributed to additional CFP lifetime quenching caused by TauRD-CFP/YFP

**Fig. 7.**
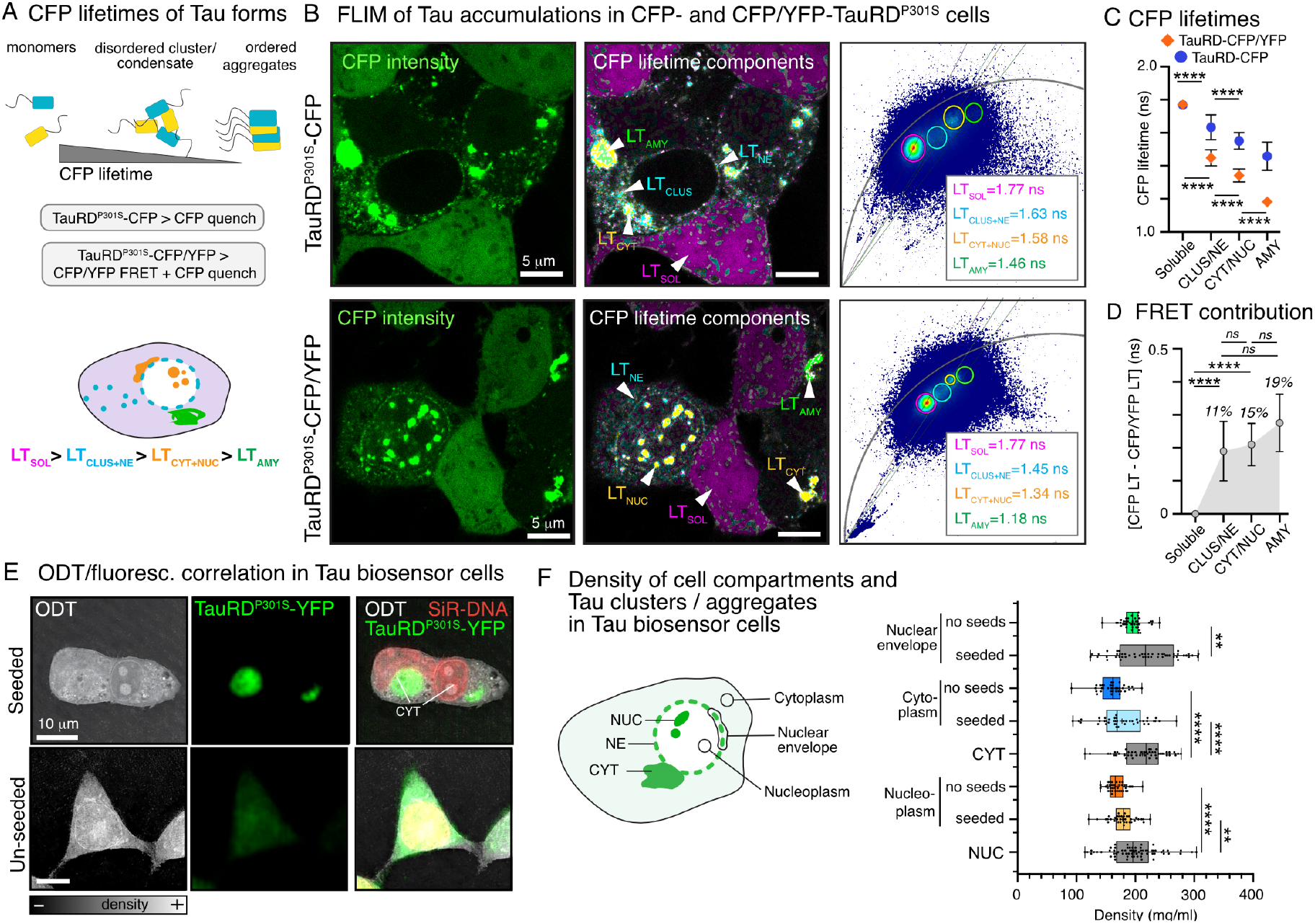
Maturation of seeded Tau assemblies in cells. **(A)** Principle of CFP lifetime FLIM in Tau biosensor cells expressing TauRD^P301S^-CFP, or TauRD^P301S^-CFP and TauRD^P301S^- YFP (TauRD^P301S^-CFP/YFP). CFP lifetime is quenched by dense molecular packing in TauRD^P301S^-CFP accumulations and by both dense packing and Tau-Tau interactions in TauRD^P301S^-CFP/YFP accumulations. **(B)** Seeded Tau biosensor cells (top: TauRD^P301S^- CFP cells; bottom: TauRD^P301S^-CFP/YFP cells). CFP intensity and CFP lifetime components superimposed on CFP intensity (fit-free defined based on ROIs in phasor plots) are shown. Lifetime components were defined for free soluble Tau (LT_SOL_, pink), Tau foci in cytosol (CLUS) and at the nuclear envelope (NE; LT_CLUS+NE_), cytosolic (CYT) and nuclear (NUC; LT_CYT+NUC_) Tau aggregates, and amyloid-like cytosolic Tau aggregates (AMY; LT_AMY_). Scale bars = 5 μm. **(C)** Lifetimes of Tau accumulation types in TauRD^P301S^- CFP and TauRD^P301S^-CFP/YFP accumulations. Data shown as mean±SD, comparison of Tau accumulation types within cells: one-way ANOVA with Tukey post-test. **(D)** FRET contribution to CFP lifetime quenching in seeded TauRD^P301S^-CFP/YFP cells, estimated by subtracting lifetimes of Tau accumulation types measured in TauRD^P301S^-CFP/YFP cells from that measured in TauRD^P301S^-CFP cells. % values give the proportion of plotted values to entire CFP lifetime quenching in TauRD^P301S^-CFP/YFP cells. Data shown as mean±SD. **(E)** Examples of ODT overlaid with correlative fluorescent images of seeded and unseeded TauRD^P301S^-CFP/YFP cells. **(L)** Quantification of densities (mg/ml) in Tau accumulations (CYT, NUC) and subcellular compartments (cytoplasm, nucleoplasm, nuclear envelope, and nucleolus) determined for ROIs defined in RI tomograms. Note, nuclear envelope density in seeded Tau biosensor cells was determined as proxy for Tau foci at the nuclear envelope. N = 15-66 measurements, box plot shows full data range (Min to Max) with all data points, line indicates median, cross indicates mean. Comparison within aggregate type and subcellular compartments: One-way ANOVA with Tukey post-test, or Student T-test for nuclear envelope.

FRET in dense Tau assemblies, in addition to higher density. To estimate the FRET contribution, as a proxy for the amount of Tau-Tau interactions, we subtracted the lifetime of Tau foci (NE and CLUS) or aggregates (CYT) in cells expressing TauRD-CFP/YFP from that in cells expressing only TauRD-CFP (Fig. 7D). This indirect analysis suggested that the contribution of FRET to the observed LT reduction in Tau aggregates (CYT ∼ 15%, AMY ∼19%) was only slightly higher (non-significant) than that in Tau foci (NE and CLUS ∼ 11%), indicating somewhat enhanced Tau-Tau interactions in Tau aggregates than in Tau foci - in line with the polymerization we observed in aging Tau/RNA/PEG condensates *in vitro*. Together, FLIM of Tau in biosensor cells suggested that early Tau foci are denser than soluble cytosolic Tau but less dense than subsequently forming larger cytosolic Tau aggregates, which also show stronger Tau-Tau interactions.

To directly measure the mass density of Tau foci and aggregates, we performed ODT on Tau biosensor cells, with or without seeding using aged Tau/RNA/PEG condensates (Fig. 7E). In the RI tomograms, based on ROIs defined in the TauRD^P301S^-YFP fluorescence channel, we measured the density of aggregates in the cytosol (CYT) and the nucleus (NUC). Since Tau foci at the nuclear envelope were too small to be outlined individually in RI tomograms, we measured the density of the nuclear envelope as a proxy for density changes induced by local Tau foci formation. In addition, we also measured subcellular density changes occurring upon seeding in the cytoplasm, the nucleoplasm, and the nucleoli of seeded and un-seeded cells (Fig. 7F). We found that the nuclear envelope in seeded biosensor cells (218 mg/ml) had a density similar to Tau/RNA/PEG condensates *in vitro*, which was significantly denser than that of un-seeded cells (196 mg/ml). Cytosolic (CYT, 212 mg/ml) and nuclear (NUC, 198 mg/ml) Tau aggregates were denser than the surrounding cytoplasm or nucleoplasm, respectively. In contrast to own previous data showing that the nucleoplasm is generally less dense than the cytoplasm (*22*), we found the density of cytosol and nucleoplasm (outside of Tau aggregates) to be similar in both seeded and un-seeded Tau biosensor cells. This can likely be accounted to the enrichment of overexpressed TauRD^P301S^-YFP in the nucleus (compare TauRD^P301S^-YFP fluorescence in nucleus versus cytoplasm in Fig. 5B) and/or to the general impairment of the nucleo- cytoplasmic transport in these cells (*11*). Together, the ODT data showed that different Tau accumulations in cells have similar densities as *in vitro* condensates. However, although lower CFP lifetime quenching by FLIM had suggested lower molecular densities for Tau foci at the nuclear envelope (NE) than for cytosolic Tau aggregates (CYT), we detected a higher density for the nuclear envelope by ODT. This apparent discrepancy could be caused by spatial imprecision in our density measurement of Tau foci at the nuclear envelope, by the contribution of nuclear envelope structures to the measured densities, and by the contribution of lipids in the nuclear envelope bilayer, which change the RI (alpha increment for lipids is lower than for proteins), hence the mass density.

In summary, the experiments in Tau biosensor cells showed that aged Tau condensates, for which we detected increased parallel arrangement by XL-MS and small, amyloid-containing Tau clusters at dense/light phase interphases, can seed Tau aggregation leading to the formation of different Tau accumulation types. initiated by small Tau. Early Tau foci in cytoplasm and at the nuclear envelope seem to have a different molecular densities and Tau arrangement than larger cytoplasmic aggregates. Whether they undergo condensate-like maturation and develop additional Tau seeds remains to be clarified.

## Discussion

In this study, we provide structural insights into Tau amyloid formation inside Tau condensates. Using optical trap viscoelasticity measurements, ODT, and crosslinking mass spectrometry, we demonstrate that condensates formed by Tau and RNA (in the presence of the molecular crowder PEG) undergo a substantial internal molecular rearrangement during their maturation, leading to the formation of condensate-spanning and shell-like elastic protein networks and amyloid-like Tau seeds. In cells, Tau condensate aging becomes relevant in the cytosol, where small condensed Tau foci appear first upon seeding, particularly around the nucleus at the cytoplasmic side of the nuclear envelope. Maturation of these Tau condensates could further accelerate intracellular Tau aggregation by producing additional Tau seeds.

### Mechanical properties

Through characterization of the complex, frequency-dependent viscoelasticity of Tau/RNA/PEG condensates *in vitro*, we find that the storage modulus (elasticity) dominated the condensate material properties over all frequencies tested and increased further with Tau/RNA/PEG condensate aging. The average storage modulus of Tau/RNA/PEG condensates increased from ∼10^2^ Pa in the first 2 h, and to ∼10^3^ Pa thereafter, and was therefore higher than that of amyloid-containing FUS hydrogels (∼10^2^ Pa; (*34*)) and similar to the elasticity of brain tissues (∼10^3^ Pa; (*35*)). In cells, the stable and “hard” structure of aged Tau condensates could provide them with a special resistance to cellular degradation systems. While proteasomal degradation is mostly restricted to the degradation of soluble proteins (*36*), oligomer assemblies of Tau (*37*) - and potentially Tau condensates (*38*) - can be degraded by autophagy. However, cellular Tau condensation appears on MTs (*39–42*), and it remains to be tested whether autophagy could degrade MT-bound, aged Tau condensates. Furthermore, maturation-related changes in the viscoelastic properties of condensed Tau on MTs could largely influence the rigidity and stability of individual MTs, and especially of Tau coated MT bundles.

The elastic dominance in Tau/RNA/PEG condensates argues against a liquid-like character and Maxwell fluid-like aging of Tau/RNA/PEG condensates, which has previously been described for different condensates (*15*, *16*). In contrast, our data suggests a progressive dynamic arrest and the formation of large networks and shells formed by polymerized Tau. Viscoelastic phase transitions have previously been observed for protein-RNA condensates (*13*), cytosolic synapsin-1 condensates (*43*) and MEC-2 stomatin *in vivo* and *in vitro* (*16*). Most similar to our Tau/RNA/PEG condensate data, an elastic behavior and internally evolving protein network structure has been reported for condensates formed by the RNA-binding protein hnRNPA1 (*13*). We also observed a strong dependence of the storage modulus on frequency for both young and mature condensates, indicating that they may not be terminally arrested but still capable of a certain degree of molecular rearrangement.

Importantly, in hnRNPA1 condensate preparations, amyloid-like hnRNPA1 fibrillization was observed outside aging hnRNPA1 condensates through recruitment of naïve molecules from the dilute phase (*13*), which we could not observe for aged Tau/RNA/PEG condensates. Instead, we observed mostly small beta-structure containing Tau foci inside and at the interface of the dense and the dilute phase of aged Tau/RNA/PEG condensates, and to a lesser extend also in Tau/PEG condensates, which correlated with the potential of the aged condensates to seed cellular Tau aggregation. The formation of small Tau species having the potential to seed Tau aggregation is in line with our previous data showing that aged Tau/RNA condensates have a higher Tau seeding activity than *in vitro*- fibrillized Tau when added to cells or to recombinant Tau (*11*).

### Molecular restructuring

Our optical trap data collection, which started a few (∼20 min) minutes after condensate formation, revealed an elastic behavior of even young Tau condensates. This indicated that “Tau/RNA/PEG liquids” had aged on the minute scale, rendering the liquid state essentially ephemeral. In agreement with this, we observed progressive hardening during a single measurement over the course of a few seconds. Moreover, we observed a 10-fold solidification over the course of 2 h, with a noticeable variability across all condensates tested. According to our ODT data, we account this to the evolution of highly heterogeneous condensates with underlying subdomains consisting of mesh-like Tau networks that give rise to spatially distinct, dense domains, eventually forming a condensate “shell”. Similarly, a liquid/solid coexistence of Tau in condensates was recently suggested (*44*), which may result in a highly variable rheological signature depending on the spatial probe positioning.

Notably, the formation of stiff, amyloid-like fibrillar Tau assemblies inside condensates should lead to a loss of their spherical shape in favor of fibrillar or amorphous structures. Since aged Tau condensates maintained their spherical shape and contained only small beta-structure containing Tau “foci”, we suggest that the internal Tau polymerization led to the formation of non-amyloid Tau networks that are stable enough to produce the observed increase in elasticity and loss of molecular mobility, notably, while preserving condensate shape. Accordingly, our FRAP data showed a rapid decrease in molecular Tau diffusion and an increase in the immobile Tau fraction of condensates over time, and micro- rheology and ODT measurements indicated the formation of a molecular “shell” around Tau/RNA/PEG condensates. Interestingly, Tau/PEG condensates formed in the absence of RNA did not show polymerized Tau networks and shells. We conclude that the observed polymerization of Tau inside aging condensates is RNA driven and largely off-path to amyloid aggregation, yet responsible for producing stable, elastic Tau condensates. Interestingly, it was recently observed that Tau/RNA aggregates (not condensates) can contain alternative Tau fibril structures, in which the fibril core is assembled from Tau C-termini and which may therefore be off-path to amyloid formation (*45*). Whether the Tau polymers we observe in condensates have a similar ultrastructure, and whether off-path polymerization of Tau in condensates can be seen as protective against Tau amyloid formation – as suggested for hnRNPA1 (*13*) – remains to be clarified.

In cells, we observed that the Tau aggregation process seeded by aged Tau/RNA/PEG condensates starts with the formation of small Tau foci in the cytoplasm and at the nuclear envelope and progresses with the formation of larger cytoplasmic aggregates near to the nucleus. Whether cellular Tau foci mature similar to Tau/RNA/PEG condensates *in vitro*, leading to hard, elastic Tau assemblies and the formation of beta-structure containing Tau seeds, remains unclear. For Tau foci at the nuclear envelope, the aging into hard structures could exert unusual local forces on the nuclear envelope that contribute to the impairment of envelope and nuclear transport integrity, as observed in AD and tauopathy brains, as well as animal and cell models of FTD (*32*, *46*).

### Tau molecular interactions

Our XL-MS data indicated, that Tau molecules undergo a partial conformational extension during the condensation process, thereby enabling Tau repeat domain interactions. This is in line with previous data from single molecule FRET measurements (*47*). In addition, we found that Tau protein-protein interactions in Tau/RNA/PEG condensates largely involved the TauRD, and that the majority of Tau molecules, based on crosslink modeling, was arranged in an anti-parallel orientation. Tau amyloid fibrils, in contrast, are built from parallel stacks of Tau repeat domains (*30*). Stably polymerized, anti-parallel Tau networks inside condensates may therefore delay amyloid formation. During condensate maturation, however, the fraction of parallel oriented Tau molecules increased, which would allow for amyloid formation by Tau repeat domains, and highly crosslinked, trypsin-resistant oligomeric Tau species formed. This is in line with our observation (here and previously (*11*)) that aged – but not fresh - Tau/RNA/PEG condensates have a high pathological seeding competence. Seeding of Tau aggregation is based on templated misfolding, in which pre-existing β-structure-rich, parallel Tau assemblies template the structural conversion of naïve, unstructured Tau molecules to adopt an amyloid fold. By amyloid-dye imaging, we observed mostly small, amyloid-containing Tau species forming inside condensates and at interphases between dense and dilute phase, which can be sufficient to seed Tau aggregation. Notably, also 24 h-old Tau/PEG condensates – showing no formation of Tau polymer networks and shells by ODT and having less amyloid seeds – increased their parallel Tau content over time (XL-MS data). This shows that parallel Tau arrangement alone is not sufficient to efficiently produce amyloid-like Tau seeds inside condensates, and that RNA (or other polyanions) is needed for this process.

To prevent the formation of seeding competent Tau oligomers in cellular condensates, which likely also contain RNA, a constant turnover and high dynamics of Tau molecules and/or efficient chaperoning mechanisms would be necessary. Since aged Tau condensates resembled porous particles into which smaller dextran molecules (5 > x > 70 kDa) could freely diffuse, it is likely that small chaperoning biomolecules and therapeutic compounds can have access to Tau molecules in the condensate interior, offering the possibility to therapeutically target the earliest stages of Tau seed formation. However, for the same reason, pathogenic Tau oligomers - having a size of a few nanometers (∼1–10 nm; (*48*, *49*)) - could also exchange between aged Tau condensates and the cytosol. Therefore, intracellular aging of Tau/RNA/PEG condensates can ferment the formation of oligomeric Tau seeds, which, when escaping from aged Tau-RNA condensates, may promote Tau oligomerization and aggregation in the cytosol.

Notably, the positively charged TauRD also catalyzes Tau’s binding to RNA, and binding of RNA strongly promotes Tau condensation (*11*, *50*, *51*), presumably by neutralizing repulsive Tau-Tau interactions. However, progressive TauRD-TauRD interactions in aging condensates appear to compete with TauRD-RNA interactions, leading to a decrease of Tau-RNA binding sites, evident from the reduction of RNA enrichment in aging condensates. RNA reduction occurred prior (or in parallel) to Tau polymerization inside condensates, and, therefore, independent of Tau amyloidogenic oligomer formation. Thus, RNA is a potent catalyzer of Tau condensation and Tau-Tau interactions but does not appear to catalyze Tau amyloid aggregation in condensates. The loss of RNA from cellular Tau condensates, could, for example, affect the wetting of Tau onto MTs, hence, MT stability (*11*).

## Materials and Methods

### Protein purification

Prior to protein production all expression plasmids were verified by Sanger sequencing. Recombinant human full-length Tau (2N4R isoform, 441 aa, ∼46 kDa) was expressed and purified as previously described (*11*, *52*). Briefly, after transformation of plasmid DNA into E. coli BL21 Star (DE3; Invitrogen), protein expression was induced at OD600 = 0.6 with 0.5 mM IPTG for ∼2 h at 37 °C. Cells were harvested, resuspended in lysis buffer (20 mM MES, 1 mM EGTA, 0.2 mM MgCl2, 1 mM PMSF, 5 mM DTT, protease inhibitors (Pierce Protease Inhibitor Tablets, EDTA-free), lysed using a French press at 1000 psi for 3 times. An initial purification was done by adjusting the salt concentration to 500 mM NaCl and boiling at 95 °C for 20 min, after which cell debris and precipitated proteins were removed by centrifugation at 50’000 g and 4°C for 40 min. The supernatant containing Tau was dialyzed against Buffer A (20 mM MES, 50 mM NaCl, 1 mM MgCl2, 1 mM EGTA, 2 mM DTT, 0.1 mM PMSF, pH 6.8), sterile filtered (0.22-µm membrane filter), run through a cation exchange column (HiTrap SP HP 5mL- 17115101 Cytiva), and eluted in gradient elution with high-salt Buffer B (20 mM MES, 1 M NaCl, 1 mM MgCl2, 1 mM EGTA, 2 mM DTT, 0.1 mM PMSF, pH 6.8). Fractions containing Tau were pooled, concentrated using spin column concentrators (Pierce Protein concentrators; 10–30 kDa MWCO, Thermo Fisher Scientific), and run through a size exclusion column (SEC, Superose 6 10/300 GL- GE17-5172-01-Cytiva). Protein was eluted in 1XPBS containing 1 mM DTT and the selected fractions containing purified monomeric Tau were pooled and concentrated as before, then aliquoted, flash frozen in liquid N_2_, and stored at −80°C. Selection of fractions in both types of chromatography was assessed by SDS-PAGE electrophoresis and Coomassie staining.

6xHis-CFP-Tau and 6xHis-YFP-Tau were expressed in the same way as full-length human Tau but E. coli lysis done in a different lysis buffer (50 mM Tris, 200 mM NaCl, 20 mM Imidazole, 10% Glycerol, 0.1% Triton X-100, 1 mM PMSF, 5 mM DTT, proteases inhibitors, pH=7.4), followed by the same French press protocol. The resulting bacterial suspension was centrifuged at 35’000 g and 4°C for 1 h and the supernatant separated using His-tag affinity chromatography with imidazole gradient elution (HisTrap column 5mL- 17524801- Cytiva; Equilibration buffer: 50 mM Tris, 200 mM NaCl, 20 mM Imidazole, 10% Glycerol, 0.1 mM PMSF, 0.5 mM TCEP, pH=7.4; Elution buffer: 50 mM Tris, 200 mM NaCl, 500 mM Imidazole, 10% Glycerol, 0.1 mM PMSF, 0.5 mM TCEP, pH=7.4). Fractions containing 6xHis-CFP/YFP-Tau were concentrated using 50 KDa MWCO centrifugal filters and further purified by SEC. Protein was eluted in 1XPBS containing 0.3 mM TCEP and fractions of interest concentrated as before, flash frozen in liquid N_2_, aliquoted, and stored at −80°C. Concentrations of Tau proteins were determined by BCA assay (Pierce).

### N^15^ labeling of Tau

To produce N^15^ Tau for XL-MS, the E. coli BL21 (DE3) growth medium was composed of: 100 mL10X M9 medium (60 g Na_2_HPO_4_, 30 g KH_2_PO_4_, 5 g NaCl, 5 g N^15^H_4_Cl, in 1 l ddH_2_O), 10 ml 100X trace elements solution (5 g EDTA, 0.83 g FeCl_3_ x6 H_2_O, 80 mg ZnCl_2_, 13 mg CuCl_2_ x2 H_2_O, 10 mg CoCl_2_ x6 H_2_O, 10 mg H_3_BO_3_, 1.6 mg MnCl_2_ x6 H_2_O, in 1 l ddH_2_O), 20 ml 20% (w/v) Glucose, 1 ml 1M MgSO_4_, 0.3 ml 1M CaCl_2_, 1 ml Biotin (1 mg/ml), 1 ml Thiamin, pH=7.5. purification was done as described for Recombinant human full-length Tau.

### Fluorescent labeling of Tau

Tau proteins were fluorescently labeled using DyLight488-NHS ester (Thermo Scientific) following the manufacturer’s instructions. The dye was dissolved in DMSO to a final concentration of 10 µg/µl, then added in a 5-fold molar excess to the protein in PBS with 1 mM DTT for 2 h at room temperature, shaking at 250 rpm. Excess dye was removed by dialysis (Pur-A-Lyzer Mini Dialysis tubes, Sigma-Aldrich; MWCO 12–14 kDa) against 1xPBS, 1 mM DTT, pH 7.4 overnight at 4°C. The labeling degree (amount of dye/ molecule protein) was determined by measuring the final protein concentration with BCA assay and correlating it to the maximum absorbance of the attached dye. For calculations see manufacturer instructions.

### Preparation of Tau/RNA/PEG condensates for imaging, FRAP, optical trap, and ODT measurements

Prior to Tau condensate formation experiments, the Tau storage buffer (PBS, 1 mM DTT) was exchanged against 25 mM HEPES, 10 mM NaCl, 1 mM DTT by dialysis. Tau/RNA/PEG condensates were prepared by mixing Tau (containing 1% of Tau- DyLight488 for fluorescent microscopy applications; diluted in 25 mM HEPES, pH 7.4, 10 mM NaCl, 1 mM DTT adding) with polyA-RNA (Sigma; 200 - 6,000 nucleotides; containing 10% Cy5-labeled polyA-RNA for RNA imaging) and 5% (w/v) PEG8000 (Promega) (Tau:RNA ratios: 10 μM Tau:10 μg/ml RNA, 25 μM Tau:25 μg/ml RNA) (*53*). Samples were incubated at room temperature for 1h, 4h, or 24h. Nuclease free H_2_O was used for all buffer preparations to prevent degradation of RNA.

For imaging, 2–3 μl Tau/RNA/PEG condensate solution were pipetted on amine-treated glass-bottom dishes (TC-treated Miltenyi, CG 1.5). To avoid evaporation of the sample during microscopy, the imaging dishes were equipped with a ddH2O soaked tissue lining the inner edges and then closed. Condensates were imaged at a confocal laser scanning microscope (Nikon LSM A1Rsi+) using a 60x oil objective.

For analysis of amyloid-containing Tau species, Tau condensate preparations were spotted onto imaging dishes and Amytracker 630 (AT630, Ebba Biotech) added in a 1:1000 dilution from the stock of dye in DMSO.

For analysis of dextran penetration into Tau/RNA/PEG condensates, fluorescently labeled dextrans (Cascade blue-labeled Dextran-10kDa and TRITC-labeled Dextran70kDa (ThermoFisher) were pre-diluted 1:100 in 25 mM HEPES, 10 mM NaCl and then added to the Tau/RNA/PEG condensate solution in a 1:10 dilution.

Co-partitioning coefficients [C_dense_/C_light_] for Tau and RNA were determined based on respective average fluorescent intensity inside versus outside of Tau/RNA/PEG condensates.

### Viscoelasticity measurements with optical trap

Our optical-tweezers setup and associated routines have been described in detail elsewhere (*16*, *54*). Briefly, we used a Sensocell optical-tweezers platform (Impetux Optics) coupled to the right port of a confocal microscope (Eclipse TI, Nikon). Optical trapping was achieved in the focal plane of a 60× water-immersion objective (NA 1.2, Nikon), and spatial manipulation was enabled by two acousto-optic deflectors. Multiple traps were generated by time-sharing a single laser at 25 kHz, and forces were measured with a light-momentum force-detection module (Sensocell, Impetux Optics).

Samples were prepared by mixing Tau/RNA/PEG condensates with polystyrene microspheres (1 µm diameter; Micromod 01-54103) at a 1:1000 dilution. Then we mounted them between a #1.5 cover glass (22x22 mm; Menzel-Gläser) and a 1 mm-thick microscope slide (Deltalab), spaced by double-sided adhesive tape featuring a 2 mm-diameter aperture. To prevent wetting, cover glasses were treated overnight with PEG-silane as previously described (*55*).

Prior to each experiment, two beads were optically trapped at 80 mW per trap. Trap stiffnesses were calculated from the slope of the linear region of a high-frequency scan routine implemented in the platform’s software (LightAce, Impetux Optics). Subsequently, beads were positioned at opposite poles of a single condensate chosen to satisfy a diameter ratio:

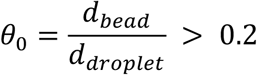

In each experiment, the left bead served as the active probe—driving condensate deformation—while the right bead was passive, reporting the mechanical response. We applied oscillatory deformations from 0.1 to 64 Hz with an amplitude of A = 0.2 µm to extract the complex spring constant of the system, (*χ_sys_*) (the two traps plus the droplet coupled to a viscous medium), as previously described (*17*):

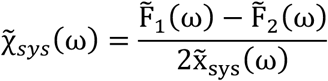

where the tilde denotes the amplitude of the Fourier transform at angular frequency *ω*. From this, we decouple the droplet’s complex spring constant χL(*ω*):

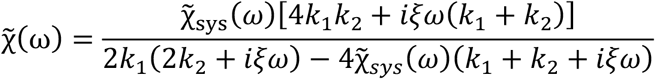

with *ξ*, the viscous drag of the droplet,

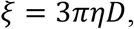

where (*η* = 1mPa s) is the viscosity of water.

To determine the static spring constant (*χ*_0_), we performed large-timescale (square-wave) oscillations with the same amplitude:

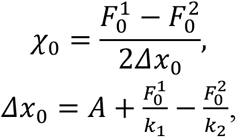

which yields the surface tension *γ* (*15*, *17*, *18*):

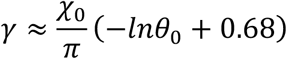

Assuming a regime of small surface tension, the complex shear modulus (*G*^O^(ω)) is given by Jawerth et al. (*56*):

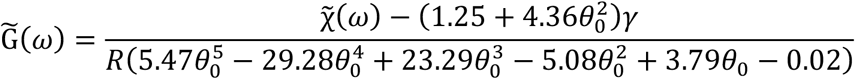

We fitted the complex modulus to a Kelvin-Voigt model weighted by the 25 percentiles to extract the elastic modulus, G_0_, and viscosity, η.

All rheology routines were programmed in LabVIEW using the manufacturer’s SDK (LightAce, Impetux Optics) and are available at https://gitlab.icfo.net/rheo/Tweezers/droplets, along with custom MATLAB scripts for computing *G*^O^(ω) from raw force data.

### Data analysis and model assumption

To quantitatively describe the frequency-dependent viscoelastic response of shell-core tau condensates, we extended the analysis beyond the classical Kelvin–Voigt (KV) model and employed a fractional Kelvin–Voigt (fKV) constitutive model. In this framework, the condensate is represented as a mechanically distinct outer shell surrounding a viscoelastic core. Because the shell component arises due to some ordering and therefore we modeled the shell as a Kelvin–Voigt element because microscopy suggested the emergence of a mechanically coherent cortical layer. The Kelvin–Voigt representation provides the simplest description of an elastic scaffold with local viscous damping, whereas the condensate interior retained clear fractional power-law rheology indicative of distributed relaxation dynamics (*20*). The classical KV model assumes a frequency-independent elastic modulus G in parallel with a Newtonian dashpot of viscosity η:

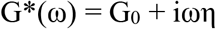

which implies:

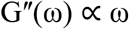

corresponding to a slope of unity in a log–log representation. This behavior reflects dissipation governed by a single characteristic relaxation mechanism. Inspection of the experimental spectra revealed substantial deviations from this prediction, particularly at early time points, where the frequency dependence of the loss modulus exhibited a significantly lower power-law exponent than expected for a classical KV material. Such sublinear scaling is inconsistent with single-timescale relaxation and instead suggests a broad spectrum of relaxation processes. We therefore modeled the core as a fractional Kelvin–Voigt model (*19*):

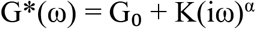

where G₀ denotes the low-frequency elastic plateau modulus, K is a fractional viscoelastic coupling coefficient with units of Pa·s^α, and 0 < α < 1 is the fractional exponent.

The complex shear modulus of the composite shell-core condensate is then is expressed as a linear combination of a standard Kelvin Voigt model and a fractional Kelvin Voigt material:

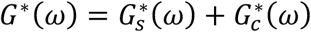

with

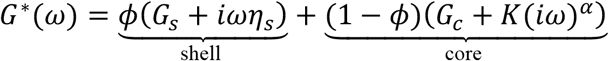

where *G_s_* and *η_s_* denote the elastic modulus and viscosity of the shell, respectively, *G_c_* is the elastic plateau modulus of the core, *K* is the fractional viscoelastic coupling coefficient, *α* is the fractional exponent, and *ϕ* represents the relative contribution of the shell to the overall mechanical response. Separating the real and imaginary components yields:

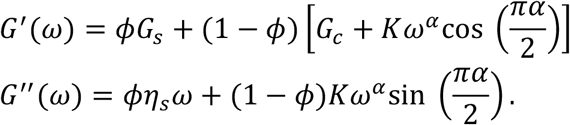

Model parameters were obtained by simultaneous nonlinear least-squares fitting of G′(ω) and G′′(ω) using a bounded Levenberg–Marquardt algorithm. As for the fractional Kelvin–Voigt model, residuals were evaluated in logarithmic space and fitting was performed simultaneously on the storage and loss moduli across the entire frequency range (minpack.lm package in R version 4.2.2).

The fractional exponent *α* was estimated from the slope of Gʺ(ω) in log–log space using linear regression and subsequently fixed during nonlinear optimization of *G* and *K*. To facilitate comparison with the classical Kelvin–Voigt model, an effective viscosity was calculated at a reference frequency of 1 Hz according to

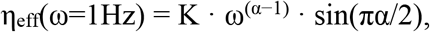

where *ω=2π rad s^−1^*. Because the fractional Kelvin–Voigt model exhibits frequency-dependent dissipation, *η_eff_* does not represent a true material constant but rather the apparent viscosity of the material at the specified frequency. Model performance was assessed using the Akaike Information Criterion (AIC), calculated from the residual sum of squares and the number of fitted parameters. Although the rheology progressively approaches the fractional Kelvin–Voigt model, our core-shell description consistently yielded similar AIC values (AIC(early)=-98; AIC(late)=-51), than the homogenous fractional KV (AIC(early)=- 93; AIC(late)=-54), model. This indicates that the core-shell decomposition and the associated meaning of the distributed viscoelastic relaxation dynamics remain significant over the whole data set despite the increased number of fitting parameters compared to the homogenous isotropic case.

The fraction *φ* can be interpreted as the thickness of the shell in relation to whole condensate.

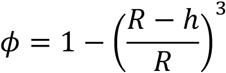

Using the derived value of *φ* and a condensate size with a diameter of 10 um, we estimate that the shell has a thickness of ∼600nm.

### FRAP of Tau condensates

Tau/RNA/PEG condensates containing ∼1% Tau-DyLight488 were imaged before and directly after bleaching with a 488 nm laser (100% intensity; 2 loops; bleach ROI diameter ∼0.5 μm for partial condensate bleaching and ∼2 μm for full condensate bleaching), and the fluorescence recovery of the bleached region was measured for 40 s. For each bleached ROI, a background ROI (outside of condensates) and a non-bleached reference-ROI (inside different condensate) of similar sizes were recorded in parallel. FRAP curves were background corrected and normalized to the background corrected reference signal. Experiments were performed on a spinning disk confocal microscope (Eclipse-Ti CSUX, Nikon) using a 60x oil objective.

### Correlative Optical Diffraction Tomography (ODT) and confocal fluorescence microscopy

The three-dimensional (3D) refractive index (RI) of Tau/RNA/PEG/PEG condensates was measured using a custom-built optical diffraction tomography (ODT) microscope employing Mach-Zehnder interferometry (*57*, *58*). Samples were illuminated with a 532 nm wavelength laser (MSL-III-532, CNI laser) and spatially modulated holograms were measured from 150 different angles, from which the complex optical fields were retrieved. By mapping the Fourier spectra of retrieved optical fields onto the surface of the Ewald sphere in the 3D Fourier space according to the Fourier diffraction theorem, 3D RI tomograms were reconstructed. Detailed principles for tomogram reconstruction can be found in (*59–61*). Image acquisition, field retrieval, and RI tomogram reconstruction were performed using custom-written MATLAB scripts (R2020a). Along with ODT images, fluorescence images or image stacks with a step size of 1 µm were acquired using a Rescan Confocal Microscope ((*62*); RCM1 from confocal.nl) equipped with a 60x water dipping objective (LUMPLFLN60XW, NA 1.0, Olympus Life Science) or 100x oil immersion objective (UPlanFl, NA 1.3, Olympus Life Science).

### ODT image analysis

For condensates, the RI of the buffer (n_buffer_ = 20mM HEPES, pH7.5, 10mM NaCl, 5% PEG) was set to a reference value of 5.5 mg/ml (= 0.5 mg/ml Tau + 0.5 μg/ml RNA in the buffer). After tomogram reconstruction (*58*, *63*), each pixel within the image was assigned a calculated RI, from which the mass density was calculated for each pixel using the following relationship: n_condensate_ = n_buffer_ + α*C_condensate_, where n_condensate_ is the average RI of the condensate, n_buffer_ the RI of the condensate suspension buffer (1.336, measured with Abbe refractometer: ABBE-2WAJ from Arcarda), α is the RI increment (0.190 ml/g for proteins and nucleic acids (*64*) and C_condensate_ the mass density (mg/ml) inside the condensates. Further details on calculating molecular mass densities from RI images can be found in (*65*). In order to calculate the mass density distribution in the condensate equatorial plane, a 0.5 mm z-projection recorded at the equatorial plane was created and the radial mass density distribution calculated using the ImageJ “radial clock scan” plug-in, which measures the average intensity in each radial scan. The provided standard deviation of these averaged radial intensities provided additional information on the variance of intensity values within each measured radial scan.

### Fluorescent lifetime microscopy (FLIM) of CFP/YFP-Tau/RNA/PEG condensates

Tau/RNA/PEG condensates formed with 10 μM eCFP-Tau and 1 μg/ml polyA-RNA in 25 mM HEPES, 1 mM DTT, pH 7.4 were imaged at 1 h or 24 h after formation. For FLIM, 2- 3 µl of each sample were pipetted on amine-treated glass-bottom dishes (TC-treated Miltenyi, CG 1.5) and imaged by confocal FLIM on a Stellaris FALCON (Leica) equipped with a 60x oil objective. FLIM of eCFP was recorded at an excitation wavelength of λEx=430 nm at laser intensity adjusted to reach a photon count of at least 50’000 counts per detected lifetime after decomposition and using a photon detector spectrum from 480 nm – 600 nm. Lifetimes for light phase (LT_light_) and dense phase (= inside condensates; LT_dense-1_, LT_dense-2_) were defined manually by placing circular ROIs of same size in phasor plots of lifetime distributions, enabling a fit-free lifetime decomposition (also see legend of Fig. S3B). Analysis was performed using LAS X software (Leica). Reduced fluorescent lifetimes of eCFP (LT_dense-1_ and LT_dense-2_) indicate a dense eCFP-Tau packing (*23*).

### Time course sedimentation assay for RNA Loss

To quantify RNA in Tau/RNA/PEG condensates during condensate aging, a time course sedimentation assay was performed. Condensates were prepared by mixing 25 μM Tau with 25 μg/ml unlabeled polyA-RNA and 5% (w/v) PEG8000 in 25 mM HEPES buffer (pH 7.4) containing 1 mM DTT and a total of 10 mM NaCl. Each reaction (10 µl) was prepared in a separate 0.5 ml Protein LoBind microcentrifuge tube (Eppendorf), each corresponding to a separate time point to avoid repeated pipetting or sample disruption. Samples were incubated at room temperature and processed at seven time points: 0, 1, 4, 6, 8, 12, and 24 h. At each time point, the respective sample was centrifuged at 16’000xg for 10 min at RT to pellet the condensates. The supernatant (2 µl) was carefully collected without disturbing the pellet. RNA concentration in the supernatant was measured using a DeNovix DS-11 spectrophotometer via absorbance at 260 nm (A260), assuming that 1.0 unit of A260 corresponds to 40 µg/ml of single-stranded RNA. Buffer-only samples were used as blanks for baseline correction. Each measurement was performed in triplicate and the experiment repeated 3 times. RNA loss from condensates was inferred from the increase in RNA concentration in the supernatant over time.

### DSS crosslinking of Tau condensates

Condensates for XL-MS experiments were prepared with a 50:50 ratio of Tau and N^15^ Tau to a final concentration of 10 μM Tau, diluted in 25 mM HEPES, 1 mM DTT with 5 mM NaCl (low salt, Tau-LS) or 300 mM NaCl (high salt, Tau-HS), pH7.4. Condensation of Tau-LS was induced with 10 μg/ml polyA-RNA (Sigma), with 5% (w/v) PEG8000, or with both polyA-RNA and PEG (*11*). Samples were incubated at room temperature for either 1 h (young) or 24 h (mature). Fibrils were also prepared with 50:50 ratio of N^14^ Tau and N^15^ Tau at a final concentration of 50 μM Tau, diluted in PBS, pH 7.4 containing 2 mM DTT. Fibrillation was induced with 0.206 mg/mL heparin (Applichem; MW = 8–25 kDa) and incubated for 7 days at 37°C. Presence of fibrils was confirmed by Thioflavin-T fluorescence, then fibrils were pelleted by ultracentrifugation at 100’000xg for 1 h at 4°C, the pellet resuspended in PBS to a final concentration of 0.5 mg/ml Tau, and sonicated at 100 W three times for 20 s. Both fibrils and condensates were cross-linked by addition of 50-fold molar excess (= 0.5 mM) disuccinimidyl suberate (DSS, ThermoScientific). Cross- linking reaction was incubated for 40 min at 37°C and 250 rpm, then the reaction was quenched by addition of Tris-HCl to final concentration 20 mM and incubated for 15 min under room temperature. Samples were inspected by bright field microscopy and SDS-PAGE, the rest stored at −20°C until XL-MS analysis.

### Crosslinking Mass Spectrometry (XL-MS) of crosslinked Tau condensates

For XL-MS analysis, crosslinked Tau condensates and soluble Tau samples were incubated in 6 M urea (8 M stock, Carl ROTH) at 37 °C for 30 min. Reduction was performed with 5 mM DTT for 30 min, followed by alkylation with 50 mM 2-chloroacetamide (CAA, Sigma-Aldrich) for another 30 min at 37 °C. Urea, DTT, and CAA were prepared in 50 mM tetraethylammonium bicarbonate (TEABC, pH 8.0). Samples were diluted to 2 M urea with 50 mM TEAB prior to enzymatic digestion. Proteins were digested overnight at 37 °C with trypsin at a 1:25 (w/w) enzyme-to-substrate ratio and Lys-C (Wako) at a 1:50 (w/w) ratio. Digested samples were acidified with 0.5% trifluoroacetic acid (TFA, pH<4, Sigma-Aldrich) and desalted using C8 cartridges (Sep-Pak Vac 1 cc, Waters). Eluates were dried in a SpeedVac (Thermo Fisher Scientific) and stored at −20 °C.

Dried samples were re-suspended in 50 µL mobile phase consisting of 30% acetonitrile (ACN, Fisher Chemical) and 0.1% trifluoroacetic acid (TFA) for size-exclusion chromatography (SEC). SEC was performed on an Agilent 1260 Infinity II system equipped with a Superdex™ 30 Increase 3.2/300 column (GE Healthcare). The flow rate was set to 0.05 mL/min with a total runtime of 60 min. Fractions were collected every 2 min, and fractions A9 to B1 were pooled and dried for subsequent LC-MS/MS analysis.

LC-MS/MS was conducted on an Orbitrap Fusion Lumos mass spectrometer (Thermo Fisher Scientific) coupled to an UltiMate 3000 ultra-high-performance liquid chromatography (UHPLC) system (Thermo Fisher Scientific), or on an Orbitrap Exploris 480 (Thermo Fisher Scientific) coupled to a Vanquish Neo UHPLC system (Thermo Fisher Scientific). Peptides were separated using an in-house packed C18 analytical column (Poroshell 120 EC-C18, 2.7 µm, Agilent Technologies) with reversed-phase chromatography. For the UltiMate 3000 system, a 160-min gradient was applied using Buffer A (0.1% formic acid (FA) in water, VWR Chemicals) and Buffer B (acetonitrile (ACN), Fisher Chemical). The gradient was programmed as follows: 0–5% Buffer B in 5 min, 5–20% in 120 min, increased to 45% in 15 min, followed by washing at 80% Buffer B at 160 min. The gradient on the Vanquish Neo system was proportionally shortened to 90 min.

Mass spectrometry analysis was performed in positive ion mode. MS^1^ scans were acquired in the Orbitrap at 120,000 resolution with a scan range of m/z 375–1600, an automatic gain control (AGC) target of 400,000, and high-field asymmetric waveform ion mobility spectrometry (FAIMS) using compensation voltages (CVs) of −40, −50, and −60. Precursors were isolated using a 1.6 m/z isolation window and fragmented by higher-energy collisional dissociation (HCD) at a normalized collision energy of 30%. MS2 spectra were acquired in the Orbitrap at 30,000 resolution with an AGC target of 100,000.

Raw files were searched against the 2N4R Tau sequence (UniProt accession number A0A024RA17) using pLink version 2.3.9 (*66*). Carbamidomethylation on cysteine (+57.0215 Da) was set as a fixed modification, and oxidation on methionine (+15.9949 Da) was defined as a variable modification. Search results were filtered at a 1% false discovery rate (FDR) at the cross-link spectrum match (CSM) level. All precursor ion intensities corresponding to identified CSMs were extracted for MS1-based quantification with in-house script in R, including monoisotopic peaks from four peak envelopes ^14^N^14^N, ^14^N^15^N, ^15^N^14^N, ^15^N^15^N. For one precursor, inter-link level, R, was calculated according to:

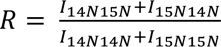

Inter-link ratios from identical residue pairs or protein–protein interaction (PPI) sites were aggregated by taking the median value of all corresponding precursors.

### AlphaLink prediction

For each cross-link, MS1-level isotope envelopes corresponding to the pure forms (^14^N^14^N or ^15^N^15^N) were required to be identified and quantified in at least one CSM. Cross-links with detectable hybrid isotope envelopes (^14^N^15^N or ^15^N^14^N) were classified as having an intermolecular contribution, whereas cross-links showing only pure-form envelopes and no detectable hybrid signal were classified as having no detectable intermolecular contribution (*27*).

The intra-residue pairs were then used as distance restraints together with the 2N4R Tau sequence for structural prediction using AlphaLink v1.0.0. The resulting intra-residue pairs were mapped onto the predicted structures, and the Cα–Cα distance was calculated for each cross-link. The radius of gyration (Rg) was calculated for residues 151–441, corresponding to the region constrained by the cross-linking data.

### Tau biosensor cell culture and seeding with condensates

HEK293 cells stably expressing the Tau repeat domain (TauRD) containing the frontotemporal dementia (FTD)-mutation P301S and fused to CFP or YFP (Tau biosensor cells; TauRD^P301S^-CFP/YFP; ATCC #CRL-3275; cells provided by Marc Diamond through Erich Wanker) were grown in 8-well imaging dishes (Ibidi). Seeding with aged Tau/RNA/PEG condensates was achieved via lipotransfection, i.e., treatment of cells with OPTI-MEM containing Tau/RNA/PEG or Tau/PEG condensates equivalent to 5 µg total Tau and 1% lipofectamine 2000 (Invitrogen) for 2 h, followed by dilution of applied condensates and lipofectamine via addition of 2.5 volumes fresh cell medium, and cells were incubated for additional 24 h under normal growth conditions.

### Imaging of Tau accumulations in Tau biosensor cells

For <u>time lapse imaging</u>, Tau biosensor cells were imaged by spinning disc confocal microscopy (TiCSU-X, Nikon) using 40x or 60x oil objectives. 10 µm z-stacks (step size 3 µm) of cells were acquired with a 40x oil objective at seven different positions each in three different wells. Images were taken in the GFP and DAPI channels for 17-18 h in 1 h intervals at 37°C overnight, starting at 6 hours after lipotransfection of aged Tau/RNA/PEG condensates into Tau biosensor cells. Quantification of aggregation types was done in max projections of the stacks in Image J. The total cell count for each recorded position and time point was measured based on DAPI+ nuclei. Tau accumulations were normalized to the respective total cell count and then averaged across all positions (n=21 positions).

STED imaging was performed on Tau biosensor cells fixed with 4% PFA in PBS at 24 h after lipotransfection, and immunostained for Lamin B1 as nuclear envelope marker using a 100x objective on a Leica Stellaris Falcon 8.

<u>Live cell FLIM</u> of CFP lifetimes was recorded in Tau biosensor cells (TauRD^P301S^-CFP and TauRD^P301S^-CFP/YFP) at 24 h after lipotransfection with aged condensates using a 60x objective on a Leica Stellaris Falcon 8. Analysis was performed using LAS X software (Leica): CFP fluorescent lifetimes were recorded at λEx=430 nm with a laser intensity of 75%. CFP lifetimes of Tau accumulation types (CLUS, NE, CYT, NUC, AMY) were determined by manually defining circular ROIs in the phasor plots leading lifetime component assignment to specific Tau accumulation types in the CFP intensity channel. Reduced fluorescent lifetimes of CFP indicated due to dense packing and, in the case of TauRD^P301S^-CFP/YFP cells, additional lifetime reduction due to CFP/YFP FRET.

<u>Optical diffraction tomograms</u> of Tau biosensor cells were obtained as described for *in vitro* Tau/RNA/PEG condensates. Nuclei were counterstained with SiR-DNA or DAPI. Alongside ODT, fluorescence images (step size of 1 µm) were acquired using a Rescan Confocal Microscope (*62*); RCM1 from confocal.nl). The mean RI value of Tau accumulations was determined by defining ROIs in the fluorescence channel and applying them to the matching RI image. For nucleoli, ROIs were defined based on RI tomograms.

### Data and statistical analysis

All data plotting, analysis, and statistical evaluation were performed using GraphPad Prism 9. Unless otherwise declared, comparison of two groups was done by Student t-test, multiple groups were compared by one-way ANOVA with Tukey or Holmes–Sidak test for multiple comparison, as indicated in the figure legends. ****P<0.0001, ***P<0.001, **P<0.01, *P<0.0.5.

### Data, code and materials availability statement

All data and code needed to evaluate and reproduce the conclusions in the paper are present in the paper and/or the Supplementary Materials. The plasmid for recombinant human Tau expression is available upon reasonable request to the corresponding author.

## Acknowledgments

External Contributions:

We thank Neus Sanfeliu-Cerdán and Frederic Català-Castro for technical help with optical trap measurements. Fluorescence microcopy and FRAP was performed at the AMBIO Charité imaging core. FLIM was performed at the imaging core of MPI- MolGen. AB and SR would like to thank Late Prof. Dr. Jochen Guck and Dr. Kyoohyun Kim for their support in establishing the ODT setup and reconstruction protocols.

## Funding

Helmholtz Association (SW)

Deutsche Forschungsgemeinschaft in the SPP2191 grant 419138680 (SW)

Deutsche Forschungsgemeinschaft in the SPP2191 grant 506373047 (SW)

Deutsche Forschungsgemeinschaft in the FOR5872 grant 545039200 (SW)

Hertie foundation grant P1200002 (SW)

Max Planck Society (SR)

Deutsche Forschungsgemeinschaft grant 528483508 – FIP 12 (SR)

## Author contribution

Conceptualization: SW, MK, FL, SR, LR, MF

Investigation: MK, MF, AB, P-L J, RS, SM, LR, A D-B, M F-C, BK N-H, JH, OK

Methodology: SW, SR, AB, MK, FL, P-L J, A D-B, M F-C, MF

Data curation: SW, MK, RS, AB, P-L J, MF

Validation: SW, RS, SM, LR, AB, M F-C, MF

Formal analysis: SW, RS, MK, AB, M F-C, P-L J, MF

Visualization: SW, MK, SM, RS, P-L J, MF

Resources: SW, TM, FL, SR, AB, P-L J, MF

Funding Acquisition: SW, FL

Project administration: SW Software: AB, M F-C, P-L J

Supervision: SW, LD, SR, MK, FL, TM

Writing—original draft: SW, MK, P-L J, MF

Writing—review & editing: SW, MK, SR, AB, FL, P-L J, SM, LR, RS, M F-C, MF

## Data, code and materials availability

All data and code are available in the main text or the supplementary materials. The plasmid for recombinant human Tau expression is available upon reasonable request to the corresponding author.

