## Supplemental Figures for "Inhomogeneous Tau polymerization, core–shell organization, and seed formation during Tau condensate aging"

**Supplementary Fig. S1**

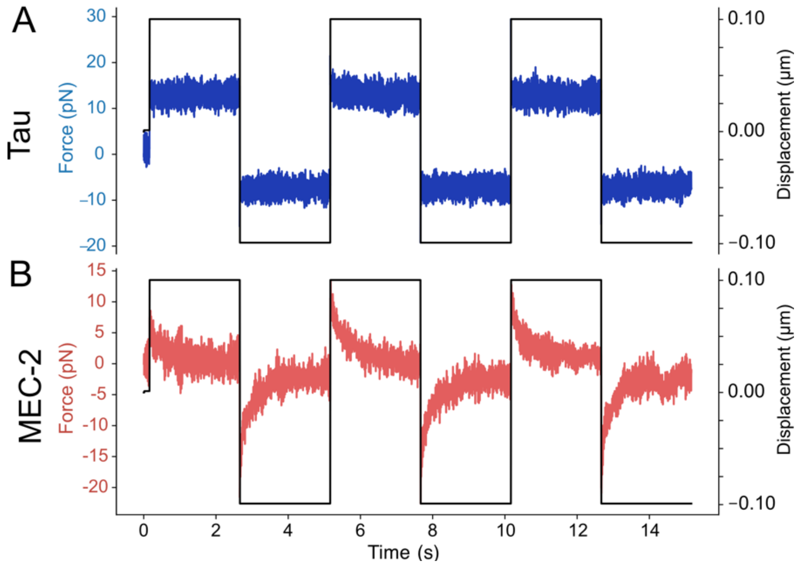


**Fig. S1. Square indentation reveals distinct viscoelastic responses of Tau/RNA/PEG and MEC-2 condensates.** Representative force responses of aged (>4 h) Tau/RNA/PEG (A, blue) and MEC-2 (B, red) condensates to square bead displacements with an amplitude of 0.2 µm. The black trace shows the imposed bead displacement. Tau/RNA/PEG condensates (A) exhibit a constant force during step indentation, consistent with purely elastic behavior. In contrast, MEC-2 condensates (B) display rapid force relaxation, indicating a rapid viscous dissipation and short timescales.

**Supplementary Fig. S2**

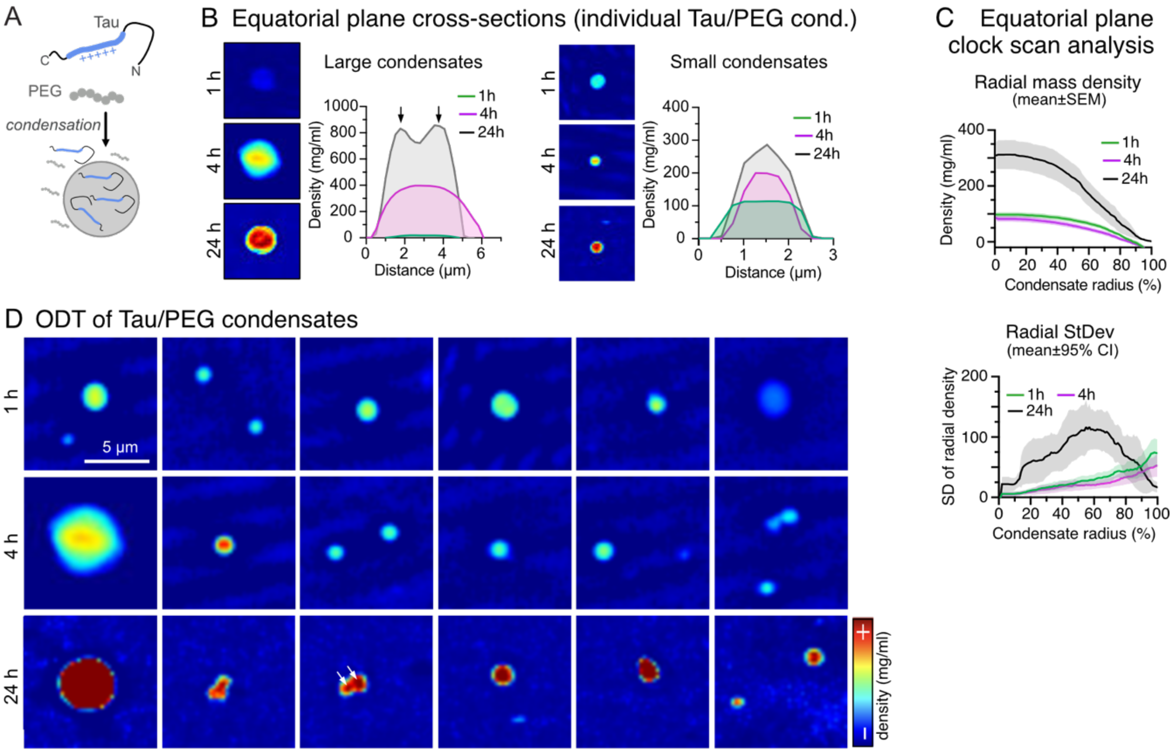


**Fig. S2. Density changes during the maturation of Tau/PEG condensates. (A)** Schematic of Tau condensation in presence of PEG**. (B)** Line profile plots through the equatorial plane of individual Tau/PEG condensates at 1h, 4 h, and 24 h. For comparison, small and large condensates were analyzed. **(C)** Average radial mass density profiles (mean±SEM) of Tau/PEG droplets at 1 h, 4 h, and 24 h. Mass density (mg/ml) is plotted from the center (R = 0%) to the border (R = 100%) of condensates (top). Radial standard deviation plot (StDev; mean±95% confidence interval; bottom) shows the average variance in density measured for each condensate at a given radius. **(D)** Gallery of equatorial plane density maps of different example 1 h, 4 h, and 24 h-old Tau/PEG condensates. Scale bar = 5 µm.

### Supplementary Fig. S3

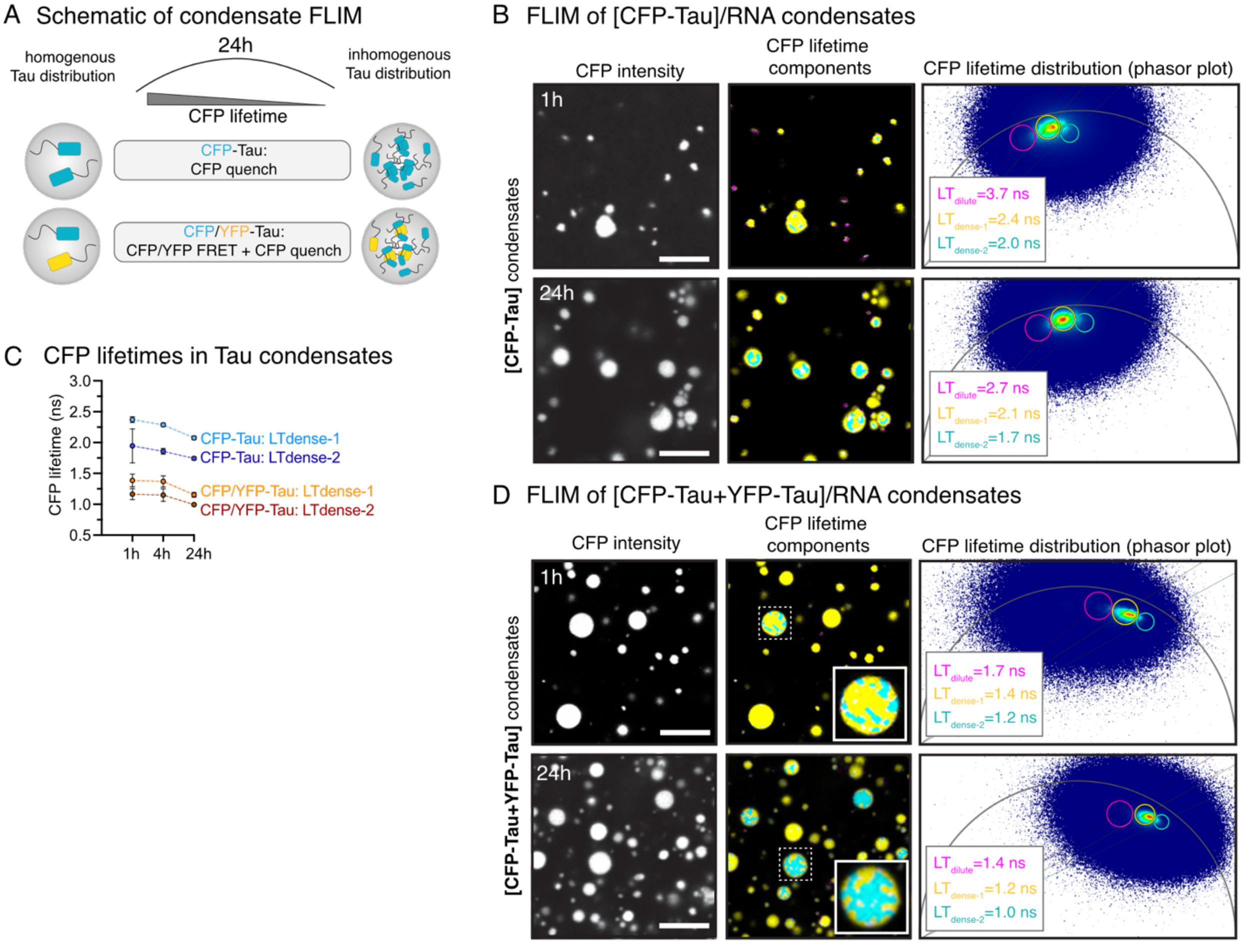


**Fig. S3. Increasing molecular density during Tau/RNA/PEG condensate aging. (A)** Principle of CFP lifetime quenching during CFP-Tau compaction during condensate maturation. **(B)** Fluorescence lifetime microscopy (FLIM) of 1 h and 24 h-old CFP-Tau/RNA/PEG condensates. Three lifetime (LT) components of decreasing LTs (dilute phase (pink, $LT_{dilute}$) > dense phase-1 (yellow, $LT_{dense\text{-}1}$) > dense phase-2 (cyan, $LT_{dense\text{-}2}$)) were fit-free assigned for individual LT-images in phasor plots of lifetime distributions. ROI sizes for LT components were kept constant between time points. ROI for $LT_{dense\text{-}1}$ was assigned to cover most pixels in condensates, and functioned as reference for positionoing the ROI $LT_{dilute}$ just left of it and the ROI $LT_{dense\text{-}2}$ to cover the pixel in the right tail of the $LT_{dense\text{-}1}$ distribution…**(C)** CFP lifetimes for LT components in condensates ($LT_{dense\text{-}1}$ and $LT_{dense\text{-}2}$) at 1 h, 4 h, and 24 h-old CFP-Tau/RNA/PEG condensates. Data shown as mean±SD, N = 8 images from 3 independent experiments per condition. **(D)** FLIM of 1 h and 24 h-old CFP/YFP-Tau/RNA/PEG condensates. LT components $LT_{dilute}$, $LT_{dense\text{-}1}$, and $LT_{dense\text{-}2}$ were fit-free assigned for individual LT-images in phasor plots of lifetime distributions.

**Supplementary Fig. S4**

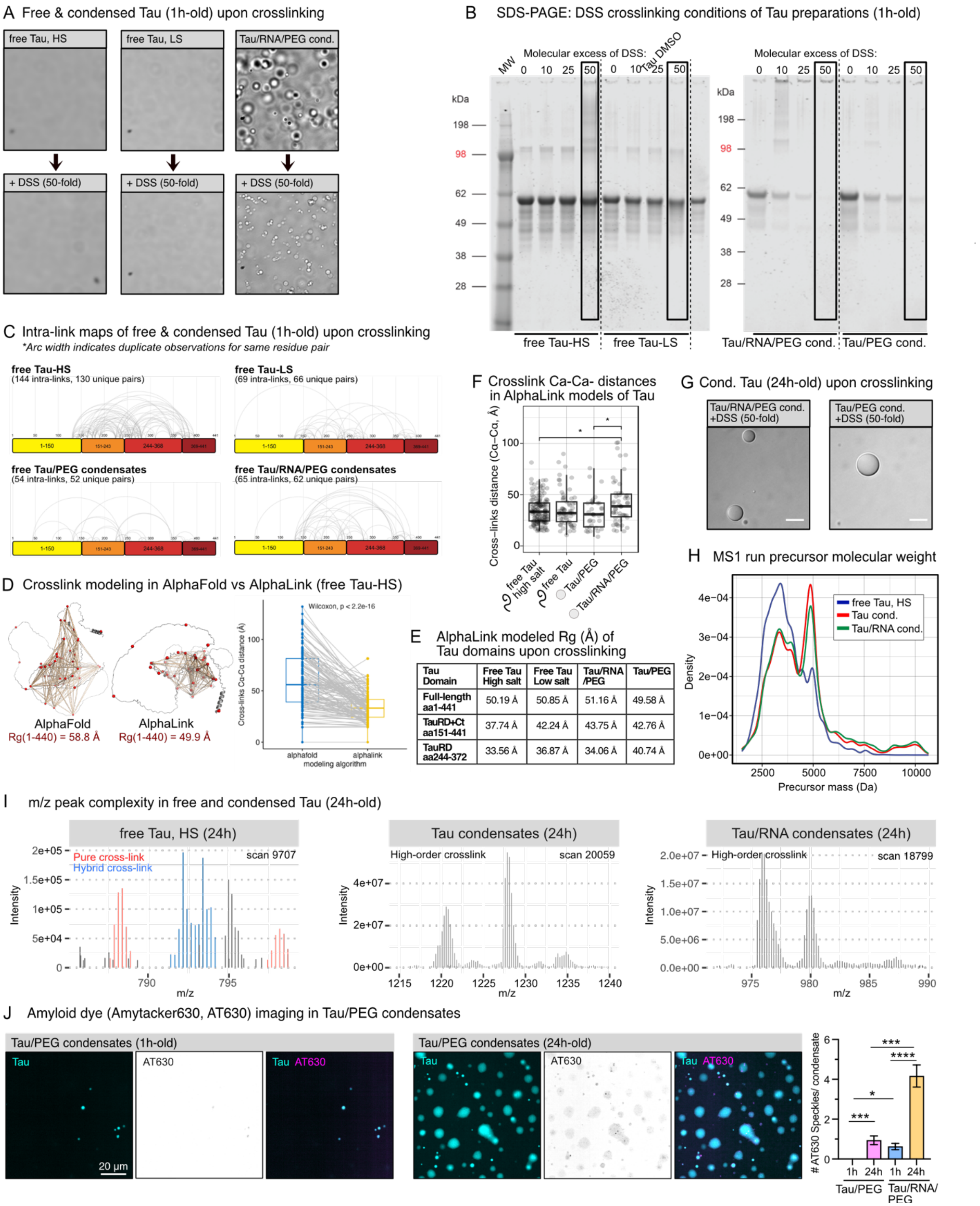


**Fig. S4. Crosslinking mass spectrometry of young and aged Tau condensates. (A)** Images of 1h-old soluble Tau in high salt (HS) and low salt (LS) buffer and of Tau condensates (incl. 5% PEG8000) before and after crosslinking with DSS. Scale bars = 20 µm. **(B)** SDS-PAGE of 1h-old Tau in HS and LS (left gel) and Tau condensates formed with PEG or without RNA (right gel) crosslinked with different molar excess (10, 20, 50-fold excess) of DSS for 40 min. **(C)** Intra-links assigned to residues in the Tau sequence (using Cytoscape) showing identified crosslinks within 1 h-old condensates and free Tau in high salt (HS) and low salt (LS) buffer. **(D)** AlphaFold versus AlphaLink predictions of Tau structure (brown ribbon) and crosslinks (red dots) in high-salt buffer. Cα–Cα distances of matched cross-links were compared between AlphaFold- and AlphaLink-predicted structures. Grey lines connect the same cross-link in both models. AlphaLink significantly reduced cross-link distances compared with AlphaFold, suggesting better satisfaction of XL-MS distance restraints. Significance was determined by paired Wilcoxon test ($p < 0.001$). **(E)** Dimensions (radius of gyration, Rg) of free and condensed full-length Tau (aa1-441), TauRD (aa244-372), and TauRD+Cterm (aa151-441) modeled by AlphaLink based on collected crosslinking data. **(F)** Analysis of distances (Å) of crosslinked Tau residues in 1h-old Tau and Tau condensate preparations. Wilcoxon's test (adjusted p-value < 0.05). **(G)** Images of 24 h-old soluble Tau in high salt (HA) and low salt (LS) buffer and of Tau condensates (incl. 5% PEG8000) before and after crosslinking with DSS. Scale bars = 20 µm. **(H)** Molecular mass (Da) of Tau species after trypsin digestion (= precursors) of 24 h-old samples. **(I)** Precursor m/z peak complexity in 24 h-old samples. Tau condensates have many species with higher order crosslinks (= high m/z ratio). **(J)** Representative images of 1 h and 24 h-old Tau/PEG condensates counterstained with Amytracker630 (AT630) to visualized amyloid-like, beta-sheet containing Tau species within condensates. Quantification of AT630+ speckles in Tau/PEG condensates, and the comparison to Tau/RNA/PEG condensates, is shown as mean±SD, One-way ANOVA with Kruskal-Wallis post-test.

**Supplementary Fig. S5**

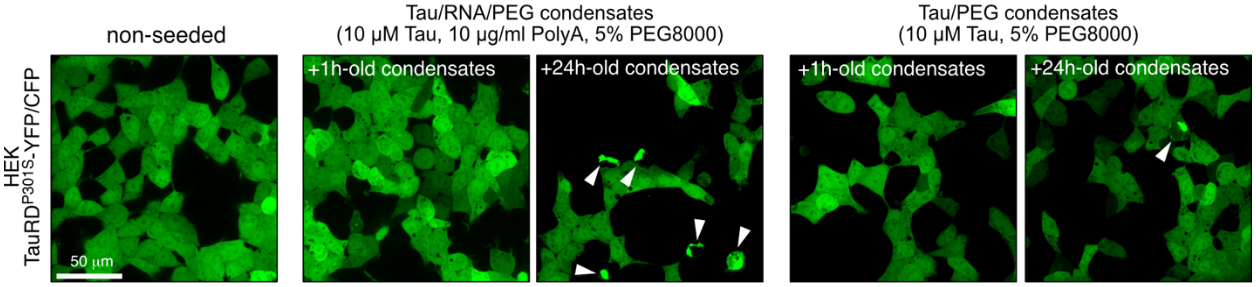


**Fig. S5. HEK TauRD$^{P301S}$-CFP/YFP cells seeded with fresh (1 h-old) or aged (24 h-old) Tau/RNA/PEG or Tau/PEG condensates.** White arrow heads indicate cytoplasmic Tau aggregates (CYT). Scale bar = 50 µm.
